# Prior vaccination enhances immune responses during SARS-CoV-2 breakthrough infection with early activation of memory T cells followed by production of potent neutralizing antibodies

**DOI:** 10.1101/2023.02.05.527215

**Authors:** Mark M. Painter, Timothy S. Johnston, Kendall A. Lundgreen, Jefferson J.S. Santos, Juliana S. Qin, Rishi R. Goel, Sokratis A. Apostolidis, Divij Mathew, Bria Fulmer, Justine C. Williams, Michelle L. McKeague, Ajinkya Pattekar, Ahmad Goode, Sean Nasta, Amy E. Baxter, Josephine R. Giles, Ashwin N. Skelly, Laura E. Felley, Maura McLaughlin, Joellen Weaver, Penn Medicine BioBank, Oliva Kuthuru, Jeanette Dougherty, Sharon Adamski, Sherea Long, Macy Kee, Cynthia Clendenin, Ricardo da Silva Antunes, Alba Grifoni, Daniela Weiskopf, Alessandro Sette, Alexander C. Huang, Daniel J. Rader, Scott E. Hensley, Paul Bates, Allison R. Greenplate, E. John Wherry

## Abstract

SARS-CoV-2 infection of vaccinated individuals is increasingly common but rarely results in severe disease, likely due to the enhanced potency and accelerated kinetics of memory immune responses. However, there have been few opportunities to rigorously study early recall responses during human viral infection. To better understand human immune memory and identify potential mediators of lasting vaccine efficacy, we used high-dimensional flow cytometry and SARS-CoV-2 antigen probes to examine immune responses in longitudinal samples from vaccinated individuals infected during the Omicron wave. These studies revealed heightened Spike-specific responses during infection of vaccinated compared to unvaccinated individuals. Spike-specific CD4 T cells and plasmablasts expanded and CD8 T cells were robustly activated during the first week. In contrast, memory B cell activation, neutralizing antibody production, and primary responses to non-Spike antigens occurred during the second week. Collectively, these data demonstrate the functionality of vaccine-primed immune memory and highlight memory T cells as rapid responders during SARS-CoV-2 infection.

## Introduction

The trajectory of the SARS-CoV-2 pandemic was fundamentally altered with the introduction of Spike-encoding mRNA vaccines in early 2021. Initial clinical trials reported greater than 90% efficacy against symptomatic infection, which has been correlated with high levels of neutralizing antibodies detected in the weeks after vaccination^1, 2^. However, these protective antibodies decline from peak levels in the months following vaccination. Coupled with the emergence of increasingly immune-evasive viral variants^3^, SARS-CoV-2 breakthrough infections, or infection of vaccinated individuals, have become increasingly common. Additional doses of mRNA vaccine are capable of boosting antibody titers and can temporarily enhance protection against symptomatic infection, though this benefit may be lost after a few months^4, 5^. Protection against hospitalization and death, however, has proved more durable following mRNA vaccination than protection against mild to moderate symptomatic infection^6, 7, 8^. Moreover, vaccinated individuals may be less likely to spread virus to close contacts^9^. Thus, protection against severe disease may be a more feasible and important goal for future vaccination strategies against SARS-CoV-2 and other rapidly evolving pathogens, such as influenza virus.

Protection against severe disease and reduced transmissibility likely result from reduced viral replication during the early stages of infection, perhaps mediated by immune memory in the form of antibodies, memory B cells, and memory T cells generated in response to prior antigen exposure. Considerable work examining the generation of immune memory following mRNA vaccination has made several key observations. Antibody titers wane from peak levels but gradually plateau over time^10, 11^. Vaccination also generates long-lasting germinal centers^12, 13^ that produce durable and stable populations of memory B cells, including cells capable of cross-binding the highly mutated RBD of Omicron variants^11, 14, 15^. Robust Spike-specific CD4 T cells are rapidly primed by the first vaccine dose and remain highly stable for months after vaccination ^14, 16, 17, 18, 19, 20^. Stable Spike-specific memory CD8 T cell populations are also generated after mRNA vaccination, and these cells are distributed across classically-defined memory T cell subsets, including stem cell memory (SCM), effector memory (EM and EM1), and terminal effector (EMRA) cells, suggesting that mRNA vaccination generates a diversity of memory T cell differentiation states^14, 18, 19, 20, 21, 22, 23^. Compared to antibodies, these T cell responses are also less susceptible to immune evasion by current variants of concern^24, 25, 26^. Memory B and T cells are capable of robust recall responses following booster vaccination, including activation of memory B cells, plasmablast expansion, production of neutralizing antibodies against Omicron variants, and activation and expansion of Spike-specific memory CD4 and CD8 T cells, supporting the functional nature of mRNA vaccine-induced immune memory^11, 20, 26, 27, 28^.

How these different components of mRNA-vaccine-induced immune memory are engaged during SARS-CoV-2 breakthrough infection, however, remains incompletely understood. Interrogating the relationships between humoral and cellular immunity could provide insights into which responses contribute to suppressing viral replication during the early stages of infection. Breakthrough infection samples collected after recovery and in convalescence show memory B cell expansion and somatic hypermutation^29, 30^ and elevated antibody titers in vaccinated individuals that experienced a recent Delta^31, 32, 33, 34^ or Omicron^29, 33, 35^ infection. However, boosting of antibodies during Omicron breakthrough infection can be negligible^21, 36^ and longitudinal analysis during breakthrough infection has suggested a delay in new antibody production^36^, placing greater emphasis on understanding the potential contribution of cellular recall responses. Although, Spike-specific T cells expand and memory CD8 T cells can become activated after breakthrough infection,^21, 30, 32, 37^ the kinetics of memory T cell and memory B cell activation relative to antibody production remain unclear, as little longitudinal data exists examining multiple components of the immune response simultaneously during the acute phase of breakthrough infection. Thus, key questions remain about the relative contribution of each component of immune memory to recall responses upon breakthrough infection. Such information may be important to guide future vaccine development.

Infectious SARS-CoV-2 virus can be recovered from nasal swabs for 7-12 days after the onset of symptoms^38, 39, 40^. Thus, components of immune memory engaged during the first week of symptoms are likely to be important for limiting viral replication and contributing to reduced disease severity. However, there remains a paucity of data measuring antigen-specific recall responses during this time frame in human viral infections, including SARS-CoV-2. Several scenarios could be envisioned. By the time symptoms are experienced, immune recall responses could already be underway. Early activation of memory B cells and plasmablasts could drive rapid production of Spike-specific antibodies, potentially clearing virus quickly enough to blunt activation of memory T cells and impede the generation of new primary immune responses against non-Spike antigens or novel variant Spike epitopes^41^. Alternatively, if pre-existing antibody titers are low and new production of antibody lags, memory T cells could play a key role in controlling viral replication during the early phase of infection, thus limiting viral spread and disease progression. It is also possible that SARS-CoV-2 possesses strategies to evade initial host immune responses enabling temporary escape from early activation of immune memory^42, 43, 44, 45^. In any of these scenarios, even a low level of pre-existing neutralizing antibodies could be an important correlate of protection from severe disease^46^. Determining which of these possibilities is observed during SARS-CoV-2 breakthrough infection will shape our understanding of the potential immunological mediators of vaccine efficacy.

In this study, we examined the kinetics of humoral and cellular Spike-specific recall responses during SARS-CoV-2 breakthrough infection of previously vaccinated individuals. We assembled two cohorts of SARS-CoV-2 infected individuals: a cohort of vaccinated individuals infected for the first time during the Omicron wave who were sampled longitudinally, and a cross-sectional cohort of vaccinated and unvaccinated individuals who tested positive for SARS-CoV-2. We measured Spike-binding and neutralizing antibody titers and the frequency, activation, and differentiation state of antigen-specific plasmablasts, memory B cells, CD4 T cells, and CD8 T cells. Collectively, these data revealed broadly accelerated and enhanced Spike-specific immune responses during infection of vaccinated individuals compared to unvaccinated individuals, whereas responses to non-Spike antigens were comparable regardless of vaccination status. Spike-specific CD4 T cells and plasmablasts expanded and CD8 T cells were robustly activated during the first week after symptom onset. In contrast, memory B cell activation and production of increasingly potent neutralizing antibodies occurred during the second week. Thus, this study highlights a coordinated immune response during SARS-CoV-2 infection of vaccinated individuals, with Spike-specific CD4 and CD8 T cells serving as early responders that may be important for mediating lasting vaccine efficacy against severe COVID-19.

## Results

### Potent neutralizing antibodies are produced during the second week of breakthrough infection

To assess the kinetics of vaccine-primed recall responses during SARS-CoV-2 breakthrough infection, we obtained peripheral blood longitudinally from a cohort of individuals who had received at least three doses of mRNA vaccines and were subsequently infected in 2022 after Omicron and related subvariants became dominant in the United States (**Fig. 1A-B**). RBD binding antibodies were detectable in all subjects prior to breakthrough infection, though titers were ∼5-fold lower than peak titers observed 2 weeks after the 3^rd^ dose of mRNA vaccine (**Fig. 1C**). Upon breakthrough infection, RBD-binding antibody titers were unchanged during the first week but increased ∼2-fold between day 7 and day 15. Binding antibody titers against the RBD from D614G and Omicron subvariants increased with similar magnitude and kinetics, suggesting that infection with Omicron drives production of circulating antibodies that retain binding to the vaccine strain as well as new variants (**Fig. 1D-E**).

**Fig. 1:**
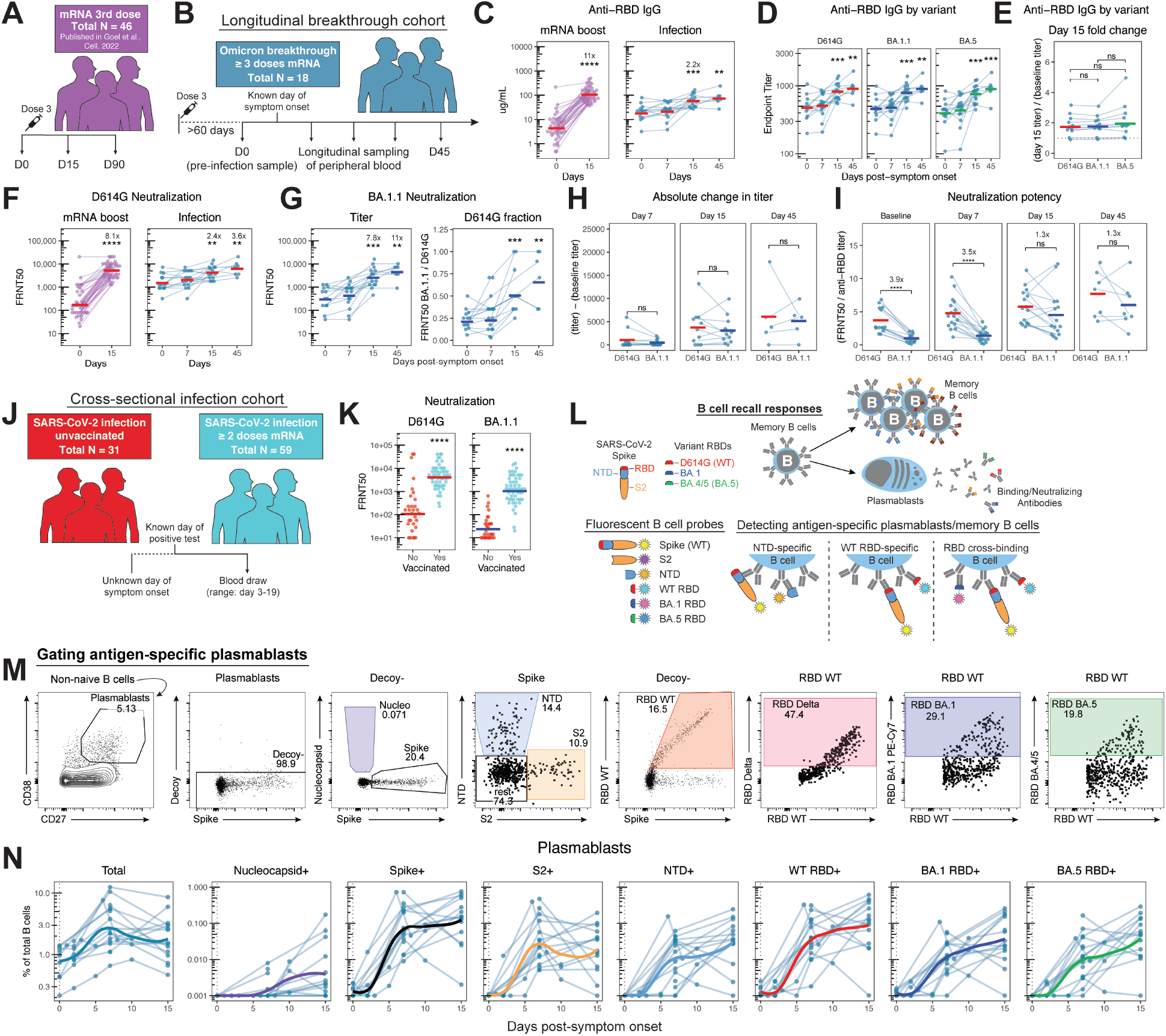
Spike-specific plasmablast expansion during the 1^st^ week of breakthrough infection precedes an increase in neutralizing antibodies during the 2^nd^ week. **A)** Schematic overview of a previously-published^11^ cohort before and after a 3^rd^ dose of SARS-CoV-2 mRNA vaccine. **B)** Schematic overview of the longitudinally-sampled Omicron breakthrough infection cohort. **C)** Wuhan-Hu-1 RBD binding antibody titers after vaccination as previously published^11^ (left) and during breakthrough infection (right). **D)** ELISA endpoint titers for RBD-binding antibodies specific for the indicated SARS-CoV-2 variants. **E)** Fold change in binding antibody titer from baseline to day 15 for each of the indicated variants. **F)** D614G neutralizing antibody titers observed after vaccination as previously published^11^ (left) and during breakthrough infection (right). **G)** Omicron BA.1.1 neutralizing antibody titers (left) and the neutralizing ratio of BA.1.1 to D614G observed during breakthrough infection (right). **H)** Change in neutralizing titers from pre-infection baseline samples calculated as absolute increase (baseline subtracted from observed) for D614G and BA.1.1. **I)** Neutralizing potency of binding antibodies calculated as the neutralizing titer divided by the paired RBD-binding titer for each variant (e.g. BA.1.1 FRNT50/ BA.1.1 binding antibody titer). Units are arbitrary. **J)** Schematic overview of the vaccinated versus unvaccinated cross-sectional cohort. **K)** D614G (left) and BA.1.1 (right) neutralizing antibody titers during infection of vaccinated and unvaccinated individuals. **L)** Schematic representation of SARS-CoV-2 Spike protein domains, B cell responses, and the flow cytometric approach for identifying antigen-specific memory B cells. **M)** Representative flow cytometric gating of total and antigen-specific plasmablasts from a single subject. All antigen-specific populations were gated on cells that did not bind a streptavidin decoy probe. Labels above plots indicate the population being displayed after upstream gating. **N)** Flow cytometric data depicting the frequency of total B cells that are plasmablasts or antigen-specific plasmablasts binding the indicated domains of Spike or Nucleocapsid during breakthrough infection. Thin lines indicate individual subjects sampled longitudinally, solid lines represent a best-fit curve of the mean response over time. Statistics calculated using Wilcoxon test with Benjamini-Hochberg correction for multiple comparisons. Statistics and fold changes without brackets are in comparison to day 0 or the unvaccinated group.

Similar results were observed for neutralizing antibody titers against the wild-type D614G strain of SARS-CoV-2 Spike. As expected, booster vaccination substantially increased D614G neutralizing titers ∼8-fold by day 15^15^ (**Fig. 1F**). During breakthrough infection, however, D614G neutralizing antibodies did not increase during the first week and rose only 2.4-fold by day 15 post-infection, with a slight additional increase by day 45 (**Fig. 1F**). Similarly, neutralizing titers against the Omicron subvariant BA.1.1 showed no quantitative increase during the first week of breakthrough infection (**Fig. 1G**). By the second week, however, BA.1.1 neutralizing antibodies increased more sharply compared to D614G, increasing 7.8-fold by day 15 (**Fig. 1G**). To test whether breakthrough infection with Omicron variant virus focused the antibody response on the variant strain, we measured the ratio of neutralizing antibody to BA.1.1 compared to D614G. Indeed, the neutralization potency of the antibody response for BA.1.1 relative to D614G significantly increased from less than 25% pre-infection to 50% by day 15 and continued to increase in some individuals (**Fig. 1G**). Notably, there was no change in this neutralization ratio during the first week of breakthrough infection. Though the fold change in BA.1.1 neutralizing antibodies was greater than that for D614G, the absolute increase in neutralizing antibodies was identical for the two variants, indicating that the antibodies produced during breakthrough infection were capable of neutralizing both BA.1.1 and D614G equivalently (**Fig. 1H**). Nevertheless, the preferential induction of antibodies capable of neutralizing BA.1.1 led to a significant increase in neutralizing potency, calculated as the neutralizing titer divided by the binding titer for each variant. Though D614G-binding antibodies had a significant 3.9-fold higher neutralizing potency than BA.1.1-binding antibodies at baseline, this deficit was largely eliminated by day 15 of breakthrough infection (**Fig. 1I**). Collectively, we observed no detectable change in circulating binding or neutralizing antibodies during the first week of Omicron breakthrough infection. Despite only modest increases in binding antibodies during the second week of infection, these antibodies were cross-reactive neutralizing antibodies against both D614G and BA.1.1, leading to significantly enhanced Omicron neutralization capacity by day 15.

We next examined how these antibody levels in previously vaccinated individuals compared to the induction of antibody following primary SARS-CoV-2 infection alone. To explore whether pre-existing vaccine-induced immunity provided an advantage for overall antibody in this setting, we measured neutralizing antibody titers at a single timepoint in a second, cross-sectional cohort of vaccinated versus unvaccinated individuals experiencing SARS-CoV-2 infection between April 2021 and December 2022 (**Fig. 1J** and **Fig. S1A**). Neutralizing antibody titers during SARS-CoV-2 infection were ∼25-fold higher in vaccinated subjects (**Fig. 1K**), suggesting that even if new antibody production is quantitatively modest and/or delayed in previously vaccinated individuals, pre-existing circulating antibodies are present at substantially higher titer. These antibodies may contribute to reducing the severity of COVID-19 in vaccinated compared to unvaccinated individuals, consistent with recent suggestions^46^.

### Spike-specific plasmablasts expand during the first week of breakthrough infection

During a recall response, memory B cells can rapidly differentiate into antibody secreting cells, including plasmablasts and plasma cells (**Fig. 1L**). To interrogate the plasmablast response during SARS-CoV-2 breakthrough infection, we used a panel of fluorescently-labelled protein tetramer probes to assess the antigen-reactivity of B cells from peripheral blood, including cells targeting SARS-CoV-2 Nucleocapsid, Spike, domains of Spike including NTD, S2, and RBD, as well as RBD variants from Delta, BA.1, and BA.4/5 (BA.5) (**Fig. 1L-M** and **Fig. S1B**). Plasmablast frequencies as a proportion of total B cells peaked at day 7 (**Figure 1N**). Plasmablasts of all Spike specificities, including those binding the RBD of BA.1 and BA.5, were detectable by day 7 and remained stable or continued to increase to day 15 (**Fig. 1N**). The temporal disconnect between plasmablast detection and increases in circulating antibody is distinct from other settings where detection of plasmablasts in the blood coincides with increased antibody titers^12, 47^. Notably, Nucleocapsid-specific plasmablasts were of low frequency overall and were rarely observed prior to day 15, when they remained 12-fold less abundant compared to Spike-specific plasmablasts (**Fig. 1N** and **Fig. S1C**). These data support the conclusion that vaccinated individuals mount rapid and robust Spike-specific plasmablast responses during SARS-CoV-2 infection, most likely arising from pre-existing Spike-specific memory B cells.

### Variant cross-binding memory B cells are activated during the second week of breakthrough infection

Rapid Spike-specific plasmablast responses in vaccinated individuals suggested a role for vaccine-primed memory B cells during breakthrough infection. The same flow cytometric approach allowed us to interrogate the activation and expansion of antigen-specific memory B cell populations (**Fig. 2A** and **Fig. S1B**). The frequencies of B cells recognizing full length Spike or individual domains of Spike were stable during the first week of breakthrough infection (**Fig. 2B** and **Fig. S1D**). In contrast, Nucleocapsid specific memory B cells were nearly undetectable in most subjects at these early time points (**Fig. 2B** and **Fig. S1D**). Spike- and Nucleocapsid-specific memory B cells expanded during the second week of infection, though the latter remained ∼80-fold lower in frequency at day 15 (**Fig. 2B**). Among Spike-specific memory B cells, S2-specific B cells did not significantly expand during the course of breakthrough infection (**Fig. 2B**). In contrast, NTD- and RBD-binding B cells significantly increased in frequency by day 15, including cells reactive with BA.1 and BA.5 variant RBDs (**Fig. 2B** and **S1D**). Memory B cells (and plasmablasts) that bound BA.1 and BA.5 without binding WT RBD were not detected at these time points in blood (**Fig. S1E-F**). This observation suggests that initial B cell recall responses during Omicron breakthrough infection were driven predominantly by vaccine-primed memory B cells that retain cross-reactivity with the vaccine strain, similar to what was observed for newly produced neutralizing antibodies.

**Fig. 2:**
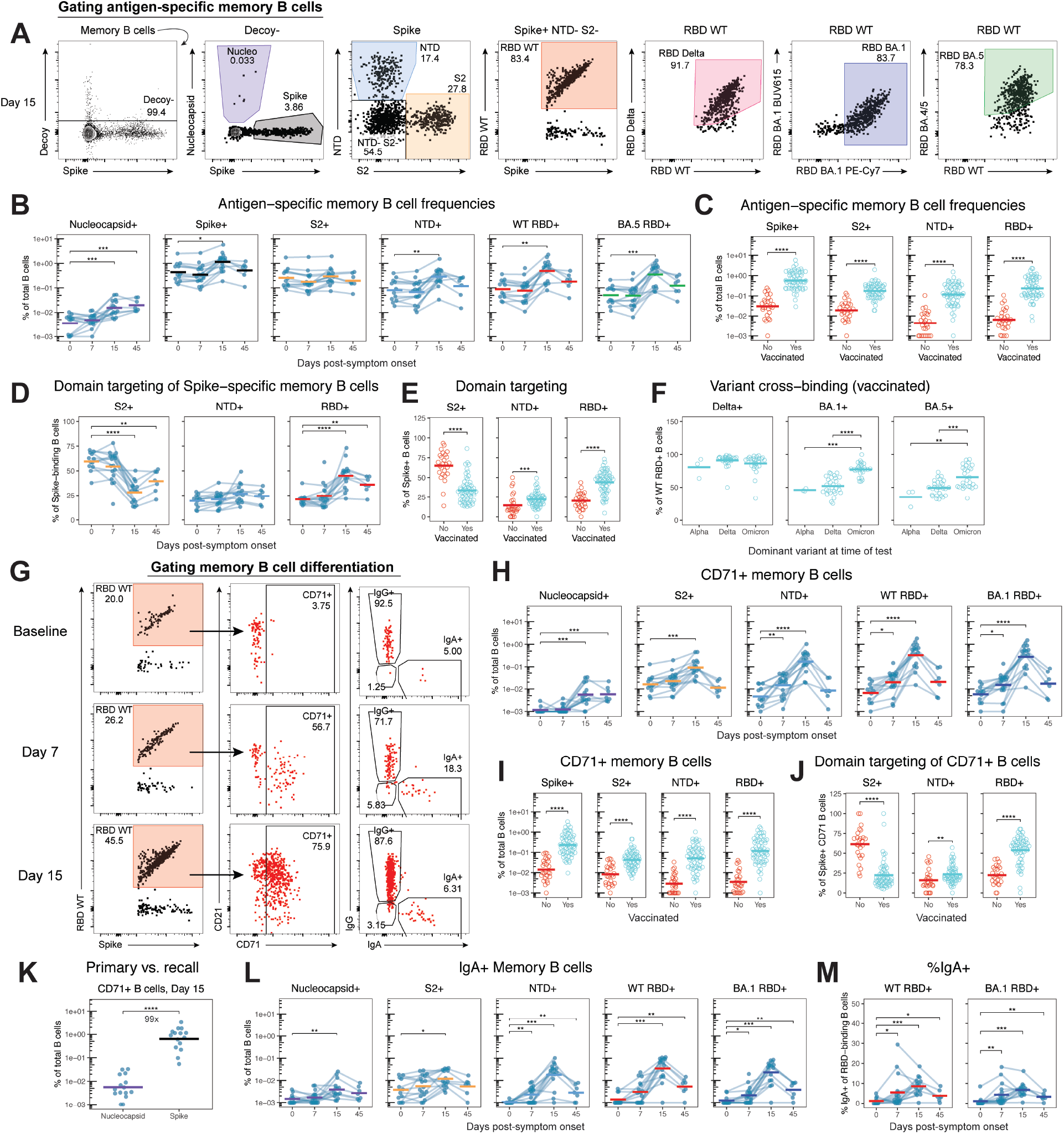
Vaccination promotes larger Spike-specific memory B cell responses that become activated, expand, and class switch during the second week of breakthrough infection. **A)** Representative flow cytometric gating of antigen-specific memory B cells from a single subject. All antigen-specific populations were gated on cells that did not bind a streptavidin decoy probe. Labels above plots indicate the population being displayed after upstream gating. **B)** Flow cytometric data depicting the frequency of total B cells that are memory B cells binding Spike or Nucleocapsid antigens during Omicron breakthrough infection. **C)** Frequency of total B cells specific for the indicated domains of Spike during SARS-CoV-2 infection in previously vaccinated or unvaccinated individuals from the cross-sectional cohort. **D-E)** Percent of Spike-binding memory B cells targeting the S2, NTD, or RBD domains during SARS-CoV-2 infection in the longitudinal cohort (D) and the cross-sectional cohort (E). **F)** Percent of WT RBD-binding memory B cells that cross bind the indicated variant RBDs during SARS-CoV-2 breakthrough infection of vaccinated individuals from the cross-sectional cohort, separated by the dominant circulating variant at the time of positive test. **G)** Representative flow cytometry plots from a single subject showing the development of activated WT RBD-specific memory B cells (CD71+) and IgA+ memory B cells during SARS-CoV-2 breakthrough infection. **H)** Frequency of total B cells that are activated (CD71+) memory B cells and bind Spike or Nucleocapsid antigens during Omicron breakthrough infection. **I)** Frequency of total B cells that are activated memory B cells and specific for the indicated domains of Spike during SARS-CoV-2 infection in previously vaccinated or unvaccinated individuals. **J)** Percent of CD71+ Spike-binding memory B cells targeting the S2, NTD, or RBD domains during SARS-CoV-2 infection. **K)** Frequency of total B cells that are CD71+ and specific for Nucleocapsid or Spike, comparing primary and recall memory B cell responses at day 15 of breakthrough infection. **L)** Frequency of total B cells that are IgA+ memory B cells and bind Spike or Nucleocapsid antigens during Omicron breakthrough infection. **M)** Percent of WT and BA.1 RBD-binding B cells that are IgA+ during breakthrough infection. Statistics calculated using Wilcoxon test with Benjamini-Hochberg correction for multiple comparisons.

The profound impact of vaccination on B cell memory was apparent in the cross-sectional cohort, which revealed significantly higher frequencies of memory B cells reacting with every domain of Spike, including S2, in vaccinated individuals (**Fig. 2C**). Vaccination did not change the frequency of Nucleocapsid-specific memory B cells detected during infection, suggesting that elevated Spike-specific memory B cell responses did not impair the ability to generate primary responses to non-Spike antigens (**Fig. S1G**). Taken together, these data demonstrate that vaccination primes durable Spike-specific memory B cells that can participate in recall responses upon breakthrough infection. Moreover, these memory B cells are present at higher frequencies than those responding to a viral antigen not contained in the vaccine.

In addition to increasing the frequency of Spike-specific memory B cells during SARS-CoV-2 infection, vaccination also altered the distribution of responses targeting different domains of the Spike protein. During the second week of infection, the proportion of Spike-specific memory B cells recognizing S2 declined significantly, whereas the proportion recognizing RBD increased, and these changes persisted at day 45 post-symptom onset (**Fig. 2D**). The amplification of RBD-specific memory B cells was associated with prior vaccination, as Spike-specific memory B cells in unvaccinated individuals predominantly targeted the S2 domain during SARS-CoV-2 infection, whereas memory B cells in vaccinated individuals were more likely to bind NTD and especially RBD, similar to previous reports of BA.1 breakthrough infection^48^ (**Fig. 2E**). Segregating individuals based on the dominant variant in circulation at the time of positive test revealed additional focusing within the RBD-specific memory B cell population. As noted previously^29^, infections during the Delta wave were associated with a modest increase in the frequency of WT RBD-binding memory B cells cross-binding Delta RBD, regardless of vaccination status (**Fig. 2F** and **Fig. S1H**). This effect was more pronounced for Omicron, as people infected during the Omicron wave had a significantly higher percentage of WT RBD-binding B cells capable of cross-binding both BA.1 and BA.5 RBD, even though many infections occurred prior to the BA.5 wave (**Fig. 2F** and **Fig. S1H-I**). Thus, vaccination led to preferential amplification of RBD- and variant cross-binding memory B cells during SARS-CoV-2 infection.

Memory B cells can also undergo phenotypic changes reflecting their activation during infection, including expression of CD71, an activation marker, and class-switch recombination to produce antibodies of different isotypes, including IgA, a key isotype for mucosal immunity^49^ (**Fig. 2G**). CD71 expression on all memory B cell populations was low prior to infection (**Fig. 2H** and **Fig. S2A-B**). Expression of this activation marker showed a modest, but statistically significant increase by day 7 after symptom onset with, on average, ∼15% of NTD- and ∼30% of RBD-specific memory B cells activated at this time point (**Fig. 2H** and **Fig. S2A-B**). Memory B cell activation was greatly increased by day 15, with 60-90% of NTD- and RBD-specific B cells expressing CD71, including memory B cells that cross-bound BA.1. and BA.5 RBD (**Fig. 2H** and **Fig. S2A-B**). Among the Spike-specific memory B cell populations, CD71 expression was lowest on S2-binding B cells and highest on RBD-binding B cells (**Fig. S2B**). Overall, vaccinated individuals had greater frequencies of activated Spike-binding memory B cells targeting all domains of Spike during breakthrough infection when compared to unvaccinated individuals (**Fig. 2I**). Thus, vaccination not only increased the total abundance of Spike-specific memory B cells, but also resulted in enhanced Spike-specific B cell activation during breakthrough infection. Similar to what was noted for overall memory B cell responses, memory B cells that were activated during infection predominantly targeted the S2 domain of Spike in unvaccinated individuals and the RBD domain of Spike in vaccinated individuals (**Fig. 2J** and **Fig. S2C**). These memory B cell recall responses also displayed a bias toward the infecting viral variant (**Fig. S2D**), though overall broad specificity remained. CD71+ memory B cells that bound Nucleocapsid were detected at day 15, but at ∼99-fold lower frequency than CD71+ Spike-binding memory B cells, highlighting the increased potency of recall responses directed against Spike compared to primary Nucleocapsid-specific responses (**Fig. 2H** and **2K** and **Fig. S2A-B**).

Examining memory B cell isotypes, there was a significant increase in NTD- and RBD-binding IgA+ memory B cells upon breakthrough infection, suggesting class switch recombination or preferential expansion of a small number of pre-existing IgA+ memory B cells (**Fig. 2L** and **Fig. S2E**). IgG+ memory B cells were also expanded at day 15 (**Fig. S2F**), but the proportion of Spike-specific B cells expressing IgA significantly increased during breakthrough infection (**Fig. 2M** and **Fig. S2G**). Furthermore, vaccinated individuals had larger populations of IgA+ B cells targeting every domain of Spike compared to unvaccinated individuals (**Fig. S2H**). Taken together, these data suggest that SARS-CoV-2 infection activated potent B cell responses in vaccinated individuals, with detection of these responses in the blood occurring predominantly during the second week after symptom onset.

### Detection of SARS-CoV-2-specific CD4 and CD8 T cell recall responses

Noting the relatively modest changes in humoral responses detected in the blood during the first week of SARS-CoV-2 breakthrough infection, we hypothesized that vaccine-primed memory T cells may play a role in early recall responses. To assess T cell responses during breakthrough infection, we performed an activation induced marker (AIM) assay^18^. Peripheral blood mononuclear cells were stimulated with megapools containing peptides derived from the SARS-CoV-2 Spike (253 peptides, 15-mers, CD4-S) or non-Spike (284 peptides, 15-20-mers, CD4-RE) proteins, and peptide-specific activation of T cells was detected by flow cytometry (**Fig. 3A** and **Fig. S3A**)^50, 51, 52^. Prior to breakthrough infection in vaccinated individuals, robust Spike-specific CD4 T cell memory was observed, and these memory CD4 T cells were predominantly Th1-like (CXCR3+ CCR6-CXCR5-) with a smaller population of circulating Tfh (cTfh, CXCR5+), consistent with previous reports (**Fig. 3B**)^14, 18^. During the first week of breakthrough infection, Spike-specific CD4 T cells expanded 2.6-fold (**Fig. 3B**). This increase was comprised predominantly of Th1 cells, though Spike-specific cTfh also trended towards an increase and expanded significantly by day 15 (**Fig. 3B**). This Spike-specific CD4 T cell response was also accompanied by a modest but significant increase in the overall frequency of activated CD4 T cells in circulation at day 7 that persisted at day 15 (**Fig. S3B-C**). Thus, upon breakthrough infection memory CD4 T cells respond with accelerated kinetics compared to humoral responses.

**Fig. 3:**
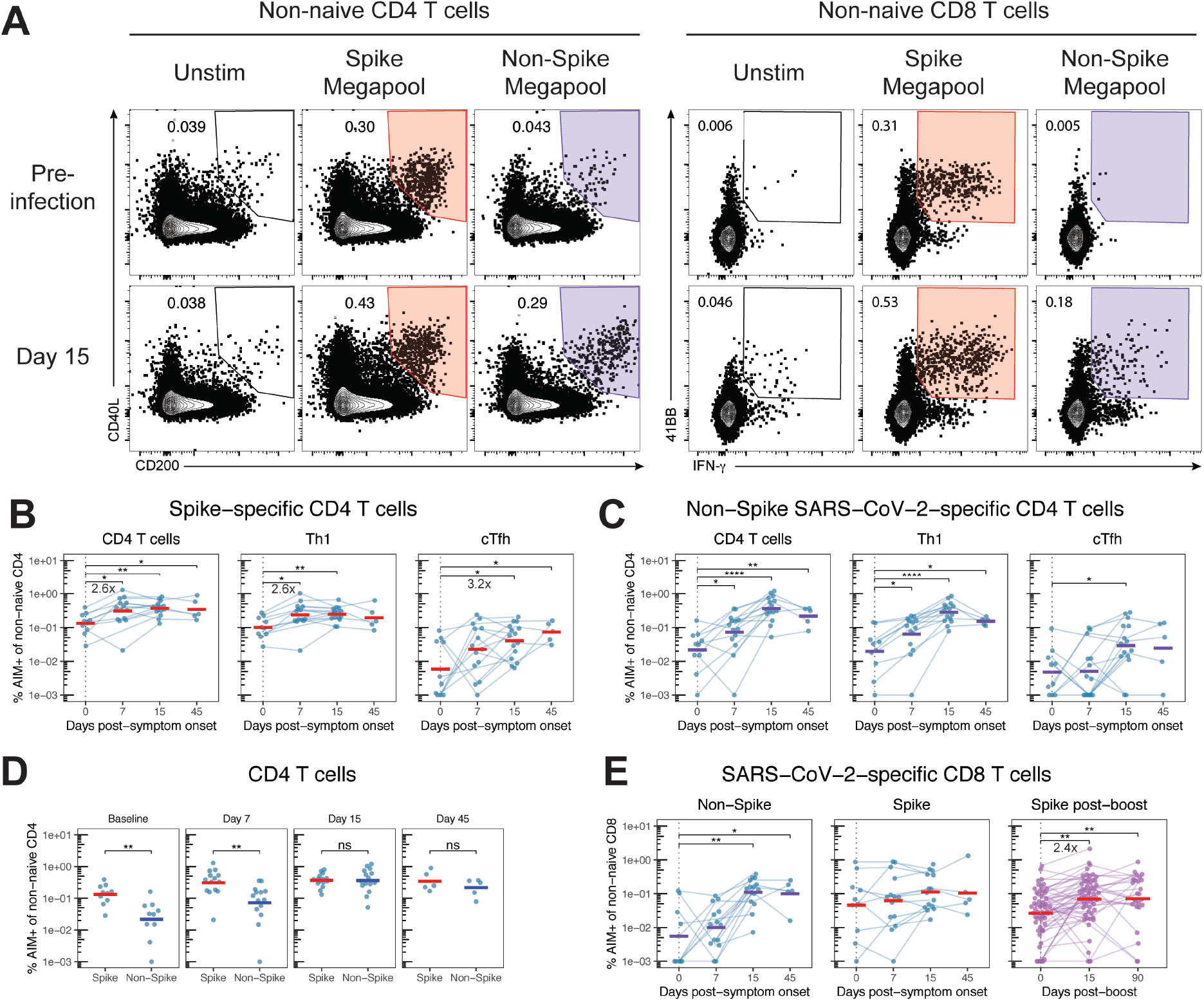
Spike-specific CD4 T cells expand during the first week of breakthrough infection while T cell responses to non-Spike SARS-CoV-2 antigens peak at day 15. **A)** Representative flow cytometric gating of CD4 (left) and CD8 (right) T cells expressing activation induced markers (AIM) after 24 hours of culture in the absence of peptides (unstim, background) or the indicated peptide megapools. **B-C)** Background-subtracted frequency of AIM+ (CD200+ CD40L+) CD4 T cells (left), Th1 cells (center, CXCR5-CXCR3+ CCR6-), and cTfh cells (right, CXCR5+) after stimulation with the Spike (B) or non-Spike (C) megapools, calculated as a percent of non-naïve CD4 T cells. **D)** Comparison of total Spike- and Non-Spike-specific CD4 T cell responses during Omicron infection. **E)** Background-subtracted frequency of AIM+ (IFNγ+ 41BB+) CD8 T cells after stimulation with the non-Spike or Spike megapools, calculated as a percent of non-naïve CD8 T cells during Omicron infection (left, center) or after a 3^rd^ dose of mRNA vaccine (right). Statistics calculated using Wilcoxon test with Benjamini-Hochberg correction for multiple comparisons.

In contrast, CD4 T cells specific for non-Spike antigens from SARS-CoV-2 were detected at low frequency prior to breakthrough infection (**Fig. 3C**). These CD4 T cell populations increased slightly by day 7 and continued to expand during the second week, reaching comparable frequencies to Spike-specific responses at day 15 (**Fig. 3C-D**). The distribution of Th1 and cTfh subsets was similar to that of Spike-specific populations (**Fig. 3C** and **Fig. S3D**)^18^. CD8 T cell responses against non-Spike antigens were similarly low-frequency prior to infection and at day 7, but were significantly expanded and equivalent to Spike-specific responses at day 15 (**Fig. 3E** and **Fig. S3E**). For a subset of samples with sufficient cell numbers, cells were stimulated with a second non-Spike megapool optimized for detecting CD8 T cell responses (621 peptides, 9-10mers, CD8-RE)^52^. CD8 T cell responses to the two non-Spike pools were highly correlated and similar kinetics were observed, though CD8 T cell responses detected with the optimized megapool were ∼2-fold greater (**Fig. S3F-H**). Thus, as previously suggested^52^, prior vaccination did not prevent the generation of primary responses to non-Spike antigens during breakthrough infection, though these responses did not peak until two weeks after symptom onset (**Fig. 3C-E**). Taken together, CD4 T cell recall responses during SARS-CoV-2 infection of vaccinated individuals were predominantly focused on rapid increases in Spike-specific responses in the first week prior to generation of primary CD4 and CD8 T cell responses against non-Spike antigens in the second week after symptom onset.

Though Spike-specific memory CD8 T cell responses were also detected prior to breakthrough infection, the frequency of Spike-specific CD8 T cells remained stable in most individuals throughout the course of infection (**Fig. 3E**). The inconsistent numerical increase in Spike-specific CD8 T cells in the blood was not due to an inability of Spike-specific memory CD8 T cells to undergo robust expansion, as these cells increased ∼2.4-fold in frequency following a 3^rd^ mRNA vaccine dose (**Fig. 3E**). Spike-specific Th1 and cTfh cells also expanded after a 3^rd^ dose of mRNA vaccine, but contrary to what was observed for CD8 T cells, the magnitude of expansion was equivalent to what was observed during the first week of breakthrough infection (**Fig. S3I** and **Fig. 3B**). This discrepancy between CD4 and CD8 T cells during breakthrough infection may derive from limitations in the AIM assay, which may have lower resolution for CD8 T cells than other approaches and only allows assessment of quantity, not activation state, of antigen-specific T cells in circulation.

### HLA class I/peptide tetramer staining reveals early activation of Spike-specific memory CD8 T cells

To further investigate the memory CD8 T cell recall response upon breakthrough infection, we developed a multiplexed HLA class I/peptide tetramer assay enabling simultaneous detection of CD8 T cells specific for SARS-CoV-2 Spike, ORF3a, Replicase, and Nucleocapsid without stimulation *ex vivo*, allowing quantitation as well as examination of differentiation and activation state (**Fig. 4A** and **Fig. S4A-B**). In a subset of study participants expressing HLA-A*02:01 or HLA-A*03:01, we assessed CD8 T cell responses using HLA class I/peptide tetramers loaded with different peptides for each antigen (**Fig. 4B-C**). All peptides from a given antigen were loaded onto tetramers with the same two fluorophores, enabling detection of a potentially diverse set of CD8 T cell populations targeting each antigen (**Fig. 4D**). Consistent with the data from the AIM assay, Spike-specific CD8 T cell frequencies detected via HLA class I/peptide tetramer binding were stable, or only increased modestly in the blood during breakthrough infection (**Fig. 4E**). CD8 T cells specific for ORF3a, Replicase, and Nucleocapsid were undetectable in most subjects prior to breakthrough infection, but became detectable during the second week (**Fig. 4E**). Despite minimal expansion in circulation, vaccinated individuals had a significantly higher frequency of Spike-specific CD8 T cells during infection compared to unvaccinated individuals in the cross-sectional cohort (**Fig. 4F**), consistent with previous priming and expansion by vaccination. In contrast, the magnitude of CD8 T cell responses to ORF3a, Replicase, and Nucleocapsid was not significantly different between vaccinated and unvaccinated subjects upon infection (**Fig. 4F**). Thus, vaccination resulted in larger Spike-specific CD8 T cell populations during subsequent SARS-CoV-2 infection without preventing the generation of *de novo* responses to non-Spike antigens.

**Fig. 4:**
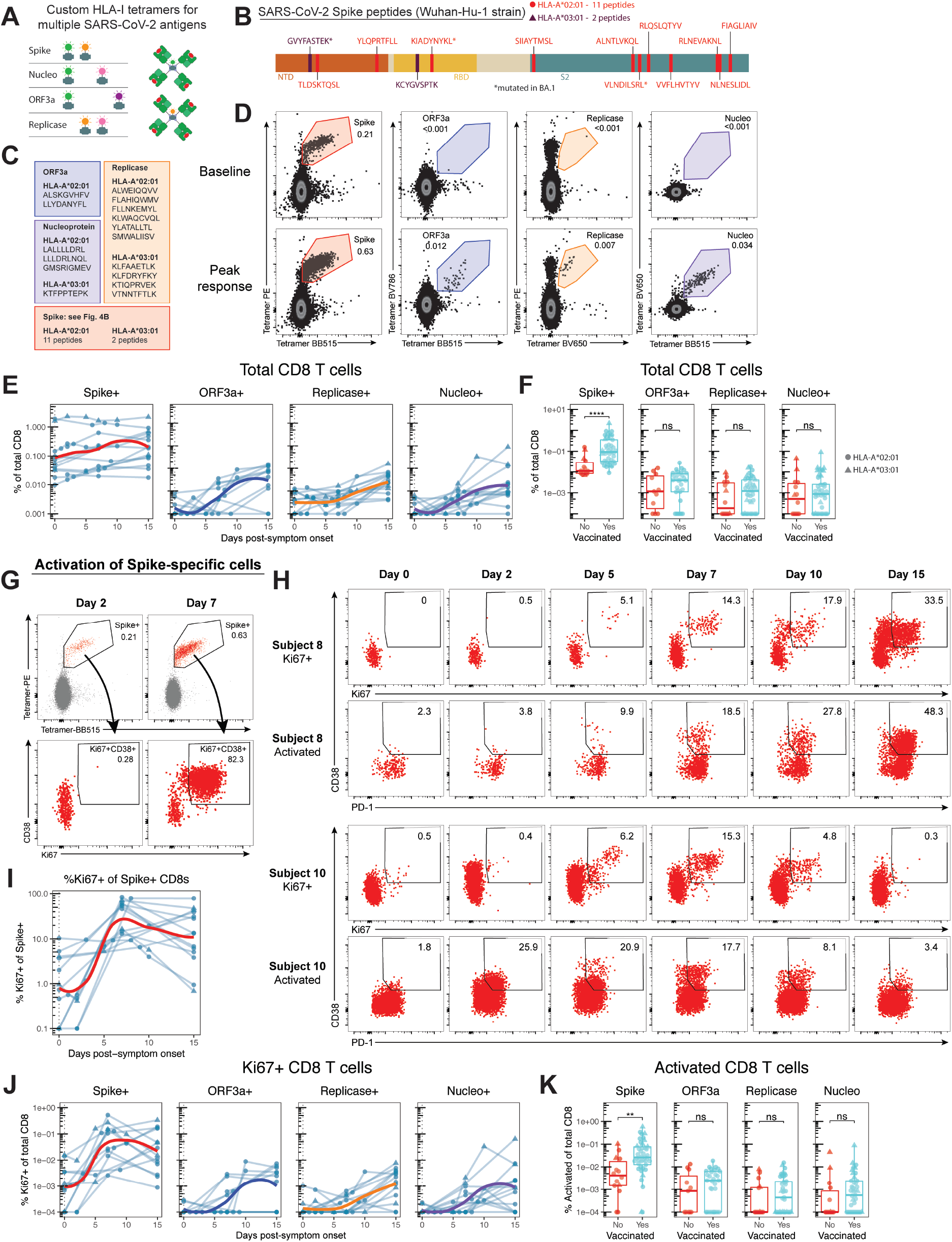
Vaccination promotes greater Spike-specific CD8 T cell responses that are activated during the first week of breakthrough infection without impeding primary responses to non-Spike antigens. **A)** Schematic representation of multiplex HLA-I/peptide tetramer assay for SARS-CoV-2 antigens using distinct combinations of fluorophore-conjugated streptavidin to assemble tetramers for each antigen. **B)** Diagram of the SARS-CoV-2 Spike protein highlighting structural domains and indicating peptides included in Spike-specific HLA-I tetramer pools for HLA-A*02:01 (red) and HLA-A*03:01 (purple). **C)** List of peptides included in HLA-I tetramer pools to identify CD8 T cells targeting SARS-CoV-2 ORF3a, Replicase, and Nucleocapsid. **D)** Representative flow cytometric gating of antigen-specific CD8 T cells identified using HLA-I tetramers with distinct dual-fluorescence profiles for each antigen. **E)** Flow cytometric data quantifying the percent of total CD8 T cells that bind HLA-I/peptide tetramers from each antigen during SARS-CoV-2 breakthrough infection. **F)** Frequency of total CD8 T cells that are specific for the indicated SARS-CoV-2 antigens during SARS-CoV-2 infection in previously vaccinated or unvaccinated individuals. **G-H)** Representative flow cytometry plots from 3 subjects depicting the activation of Spike-specific CD8 T cells during SARS-CoV-2 breakthrough infection, highlighting peak activation (G) and activation kinetics (H) of Spike-specific CD8 T cells in red. **I)** Percent of Spike-specific CD8 T cells that express Ki67 and at least one other activation marker (Fig. S3C-D). **J)** Percent of total CD8 T cells that are antigen-specific and express Ki67 and at least one other activation marker. **K)** Percent of total CD8 T cells that are antigen-specific and express at least two activation markers during SARS-CoV-2 infection in previously vaccinated or unvaccinated individuals (Fig. S3C-D). Statistics calculated using Wilcoxon test with Benjamini-Hochberg correction for multiple comparisons.

Given the elevated frequency of Spike-specific CD8 T cells in vaccinated individuals, we next sought to examine whether these cells showed evidence of activation and/or differentiation into effector CD8 T cells upon breakthrough infection. To this end, we employed a panel of five differentiation markers (Ki67, CD38, HLA-DR, PD-1, and CD39) to identify CD8 T cells that were activated (expressing at least 2 of 5 markers) or activated and proliferating (expressing Ki67 and at least 1 of 4 additional markers) (**Fig. S4C-D**). This analysis revealed substantial activation of Spike-specific CD8 T cells during the first week of infection, with activation beginning by day 5 in many individuals and peaking in most subjects by day 7 (**Fig. 4G-J** and **Fig. S4E**). Although expression of Ki67 was not apparent in Spike-specific CD8 T cells prior to day 5 after symptom onset (**Fig. 4H-I**), there were signs of activation as early as day 2 in some individuals (**Fig. 4H** and **Fig. S4E**).

In contrast to the early activation of Spike-specific memory CD8 T cells primed by prior vaccination, activation of CD8 T cells specific for other SARS-CoV-2 antigens was delayed, with levels of activation typically highest at day 15 (**Fig. 4J** and **Fig. S4E**). The frequency of activated CD8 T cells specific for non-Spike antigens was also substantially lower than for Spike, suggesting that vaccine-primed memory CD8 T cell responses to SARS-CoV-2 are both accelerated in activation and greater in magnitude than primary responses (**Fig. 4J**). These observations were supported in the cross-sectional cohort, where prior vaccination was associated with a significantly higher frequency of activated Spike-specific CD8 T cells with no significant differences in the magnitude of activated CD8 T cell responses directed against other antigens (**Fig. 4K** and **Fig. S4F**). Activation of total CD8 T cells in the blood was also observed at day 7 and day 15, and this activation of total CD8 T cells and SARS-CoV-2 specific CD8 T cells returned to baseline levels by day 45 (**Fig. S4G**). By day 45 there were no significant alterations in the frequencies of a wide range of CD4 and CD8 T cell populations after SARS-CoV-2 infection, suggesting that SARS-CoV-2 infection acutely activated antigen-specific memory T cell responses without permanently altering the broader T cell landscape (**Fig. S4G**). In summary, memory CD8 T cells were activated during the first week of breakthrough infection, preceding substantial increases in antibody and the induction of primary B and T cell responses to non-Spike antigens, and these activated Spike-specific CD8 T cell responses were also greater in magnitude in vaccinated individuals.

### Central memory cells dominate the circulating CD8 T cell response during breakthrough infection

Memory CD8 T cells can be categorized into subsets based on surface marker expression. Subsets including stem cell memory (SCM), central memory (CM), effector memory (EM) and terminal effector (EMRA) cells exist along a spectrum of increasing effector function and decreasing proliferative potential^53, 54, 55^. To better understand the dynamics of Spike-specific memory CD8 T cell recall responses during SARS-CoV-2 breakthrough infection, we identified naïve and seven non-naïve memory CD8 T cell subsets based on expression of 16 differentiation, migration and effector proteins in flow cytometric data (**Fig. 5A-B** and **Fig. S5A**). An unbiased dimensionality reduction of total CD8 T cells based on expression of these proteins identified two major groups of non-naïve cells: an EM lineage positive for both Granzyme B (GZMB) and CX3CR1, and a CM lineage negative for both proteins (**Fig. 5B** and **Fig. S5A-B**). The EM lineage could be further divided based on expression of CD45RA and CD27 into classically-defined EM, EMRA, and intermediate EM1 and “EM1-RA” cells. The CM lineage could be divided based on KLRG1 and CD45RA into SCM, CM, and KLRG1 expressing “CM-KLRG1” cells (**Fig. 5A**). These subsets were found in distinct regions of UMAP space separated within and between the EM and CM lineages (**Fig. 5B** and **Fig. S5A-B**). Compared to the CM lineage, the EM lineage was broadly associated with higher expression of the effector-associated proteins GZMB, CX3CR1, KLRG1, and Tbet and lower expression of the memory-associated proteins TCF1 and CD127 (**Fig. S5B**). Within the EM lineage, however, EM1 and EM1-RA cells had relatively higher expression of TCF1 and lower expression of Tbet, whereas within the CM lineage, CM-KLRG1 had higher Tbet and lower TCF1, suggesting that these may represent intermediate states that share properties with traditionally-defined subsets (**Figure S5A-B**).

**Fig. 5:**
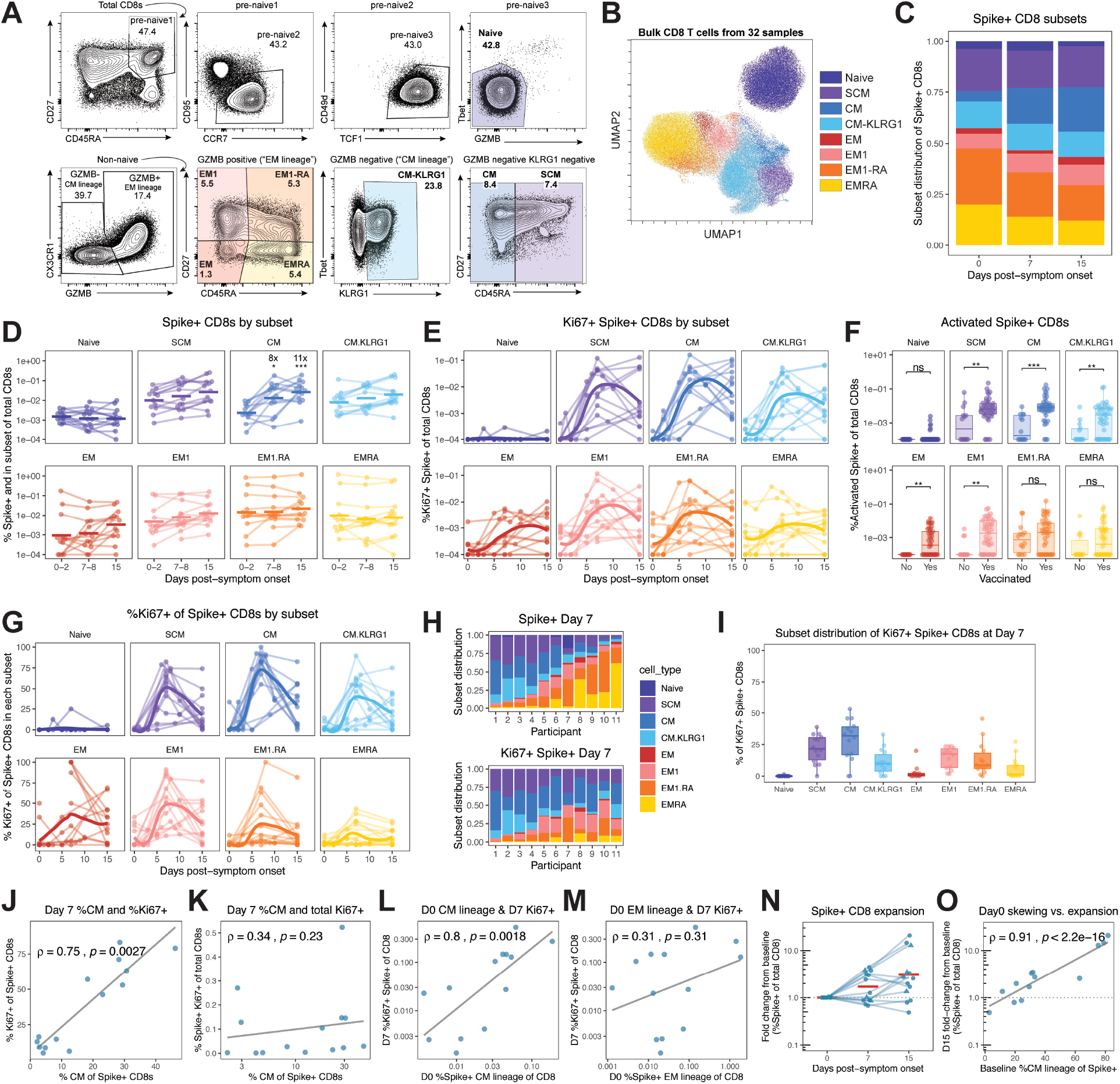
Vaccination promotes activation of central memory CD8 T cells that peaks during the first week of breakthrough infection. **A)** Representative flow cytometric gating of CD8 T cell subsets from bulk CD8 T cells. Numbers in gates represent the percent of total CD8 T cells falling in the gate. Bolded populations are represented in subsequent figures. **B)** Unbiased dimensionality reduction by Uniform Manifold Approximation and Projection (UMAP) of 16 parameters (see Fig. S4A) from flow cytometric analysis of bulk CD8 T cells from 32 unique samples. Cells are colored by manually gated subsets defined as in A. **C)** Average distribution of Spike-specific CD8 T cells into manually gated subsets during breakthrough infection. **D)** The abundance of Spike-specific CD8 T cells in each subset during breakthrough infection, calculated as the percent of total CD8 T cells. Statistics compare later timepoints to the baseline timepoint. **E)** The magnitude of the Ki67+ Spike-specific CD8 T cell response in each subset during breakthrough infection as a percent of total CD8 T cells, where cells express Ki67 and at least one other activation marker. **F)** The magnitude of the activated Spike-specific CD8 T cell response in each subset during SARS-CoV-2 infection of previously vaccinated or unvaccinated individuals, calculated as a percent of total CD8 T cells. Activated cells express at least two activation markers. **G)** The percent Ki67 positivity of Spike-specific CD8 T cells in each subset, where cells express Ki67 and at least one other activation marker. **H)** The subset distribution of total (top) and Ki67+ (bottom) Spike-specific CD8 T cells at day 7 post-symptom onset for eleven participants with paired pre-infection samples. **I)** Subset distribution of Ki67+ Spike-specific CD8 T cells at day 7 post-symptom onset. **J-M)** Correlations within Spike-specific CD8 T cell responses. J) Day 7 percent CM of Spike+ cells and day 7 percent Ki67+ of Spike+ cells, **K)** day 7 percent CM of Spike+ cells and day 7 total Ki67+, **L)** day 0 total CM lineage and day 7 total Ki67+, M) day 0 total EM lineage and day 7 total Ki67+. Statistics calculated using Wilcoxon test with Benjamini-Hochberg correction for multiple comparisons.

Prior to breakthrough infection, mRNA vaccine-induced Spike-specific CD8 T cells were predominantly distributed among EM1-RA, SCM, CM-KLRG1, EMRA, and to a lesser extent CM subsets with considerable variability among participants (**Fig. 5C-D** and **Fig. S5C**). Breakthrough infection was associated with a shift towards a higher proportion of CM cells within the overall Spike-specific CD8 T cell population (**Fig. 5C**). Though most subsets remained stable in frequency during the course of infection, the overall abundance of Spike-specific CM cells increased ∼8-fold by day 7 and this population continued to expand to day 15 (**Fig. 5D**). During SARS-CoV-2 infection, vaccinated individuals had significantly higher frequencies of these expanding Spike-specific CM cells, but also SCM, CM-KLRG1, EM, and EM1 cells compared to unvaccinated individuals (**Fig. S5D**). In summary, prior vaccination promoted enhanced CD8 T cell responses in many memory subsets during SARS-CoV-2 breakthrough infection marked by preferential expansion of CM cells.

To further interrogate how each of these vaccine-primed memory CD8 T cell populations participated in the recall response upon SARS-CoV-2 infection, we analyzed the abundance of circulating activated Spike-specific CD8 T cells in each subset. By day 7 post-symptom onset, increased activation and Ki67 expression were observed in Spike-specific CD8 T cells of all non-naïve subsets, including those generally considered to be quiescent memory CD8 T cells, with CM and SCM cells mounting the most robust responses (**Fig. 5E** and **Fig. S5E**). Large populations of activated EM1 and EM1-RA cells were also observed, whereas activated EM and EMRA cells were less prevalent in circulation. Potent activation of Spike-specific CD8 T cells across multiple memory subsets required prior vaccination, as infection in unvaccinated individuals generated significantly fewer activated, SARS-CoV-2 Spike-specific CD8 T cells among these CM, SCM, CM-KLRG1, EM, and EM1 subsets (**Fig. 5F**).

Although activated Spike-specific memory CD8 T cells were detected in all subsets on day 7 post-symptom onset, the observation that only CM cells significantly expanded provoked the hypothesis that intrinsic functional differences between subsets may influence recall responses. Indeed, the majority of CM cells were activated at day 7 in almost all subjects, whereas, on average, less than 25% of circulating Spike-specific EMRA cells were activated at any time point (**Fig. 5G** and **Fig. S5F**). SCM, CM-KLRG1, and EM1 cells were also activated in most subjects. EM1-RA cells were highly activated in some participants but minimally activated in others (**Fig. 5G** and **Fig. S5F**). The dominance of CM and SCM cells among activated Spike-specific populations was observed even in participants where these subsets represented a minority of the overall Spike-specific memory CD8 T cells detected in circulation pre-infection and on day 7 (**Fig. 5H** and **Fig. S5C**). Overall, CM cells were the most abundant subset among activated and Ki67+ CD8 T cells during breakthrough infection, with activated SCM, EM1, CM-KLRG1, and EM1-RA cells also contributing to the response (**Fig. 5I** and **Fig. S5G**). Taken together, these data demonstrate robust activation and expansion of many Spike-specific CD8 T cell subsets, especially CM cells, during the first week of SARS-CoV-2 breakthrough infection that was significantly enhanced by prior vaccination.

Though activation of Spike-specific CD8 T cells was universally observed during SARS-CoV-2 breakthrough infection, there was substantial heterogeneity in both the overall magnitude of the activated response and the proportion of Spike-specific memory CD8 T cells that showed signs of activation. We speculated that intrinsic differences in the Spike-specific memory CD8 T cell pool may partially explain this heterogeneity. Indeed, the proportion of Spike-specific CD8 T cells that were Ki67+ at day 7 was strongly correlated with the proportion of Spike-specific CM cells at day 7 (**Fig. 5J**), suggesting either that Ki67+ cells preferentially differentiate into CM cells or that CM cells are more readily able to become Ki67+ upon breakthrough infection. Despite this association, the fraction of Spike-specific CD8 T cells that were CM at day 7 was not correlated with the overall frequency of Ki67+ Spike-specific CD8 T cells (**Fig. 5K**). Instead, the frequency of Ki67+ Spike-specific CD8 T cells was strongly correlated with simply the overall frequency of Spike-specific CD8 T cells, including CM cells (**Fig. S5H**). Thus, a larger overall response, even if proportionally skewed towards less responsive EM populations, correlates with a more robust pool of activated cells. Nevertheless, we hypothesized that the pre-infection abundance of Spike-specific memory CD8 T cells in the CM lineage may better predict the magnitude of the activated response at day 7 than the abundance of Spike-specific memory CD8 T cells in the EM lineage. Indeed, there was a significant correlation between the baseline frequency of CM lineage cells and the magnitude of the Ki67+ response at day 7, whereas no such correlation existed for Spike-specific memory CD8 T cells in the EM lineage (**Fig. 5L-M**). This relationship was true despite the fact that the largest EM lineage responses were approximately 10-fold greater in magnitude than the largest CM lineage responses. These findings do not exclude the possibility that expanded populations of EM lineage cells may preferentially accumulate at the site of infection upon activation, precluding their detection in circulation, as transient activation of EM1-RA and EMRA cells was observed in a subset of individuals at day 2 (**Fig. S5E**). Though overall expansion of circulating Spike-specific CD8 T cells was not consistently observed during breakthrough infection, marked expansion did occur in some participants (**Fig. 5N**). Given the relevance of the CM lineage for Spike-specific CD8 T cell activation, we asked whether the heterogeneity in CD8 T cell expansion was associated with skewing towards the CM lineage. Indeed, there was a strong positive correlation between skewing of pre-infection Spike-specific memory CD8 T cells towards the CM lineage and the fold expansion of Spike-specific CD8 T cells at day 15 of breakthrough infection (**Fig. 5O**). Collectively, these data suggest that pre-existing Spike-specific CD8 T cells in the CM lineage are poised to drive expansion in response to antigen and highlight these CM cells as primary contributors to the circulating CD8 T cell response during the first week of SARS-CoV-2 breakthrough infection.

### Different components of immune memory respond with distinct kinetics during SARS-CoV-2 breakthrough infection

Overall, these data highlighted the distinct kinetics of recall responses for different aspects of vaccine-induced SARS-CoV-2 adaptive immunity during breakthrough infection of vaccinated individuals. To further explore this possibility, we compared when each branch of adaptive immune memory reached peak responses during the course of SARS-CoV-2 breakthrough infection. Spike-specific plasmablasts, likely derived from vaccine-primed memory B cells, were detectable within the first week of infection, but antibody titers in blood, especially variant neutralizing titers, did not increase significantly until at least day 15 (**Fig. 6**). Similarly, memory B cell activation was highest at day 15 in the blood, though it is likely that earlier activation of these cells occurs in lymphoid tissues considering the presence plasmablasts in blood at day 7. Likewise, *de novo* B and T cell responses to non-Spike antigens were only observed in the second week after symptom onset (**Fig. 6**). Alternatively, activation of Spike-specific memory CD8 T cells and significant expansion of Spike-specific memory CD4 T cells were observed by day 7 (**Fig. 6**). Furthermore, pre-existing antibody, even if not quantitatively increased during the first week, was substantially elevated in vaccinated compared to unvaccinated individuals. Thus, these data support the conclusion that the predominant systemic adaptive immune response during the first week of SARS-CoV-2 breakthrough infection consists of the activation of memory T cells, generation of new plasmablasts and the presence of pre-existing antibodies. In summary, this study highlights the broad benefits of vaccination to establish immune memory capable of accelerated and enhanced recall responses during SARS-CoV-2 infection.

**Fig. 6:**
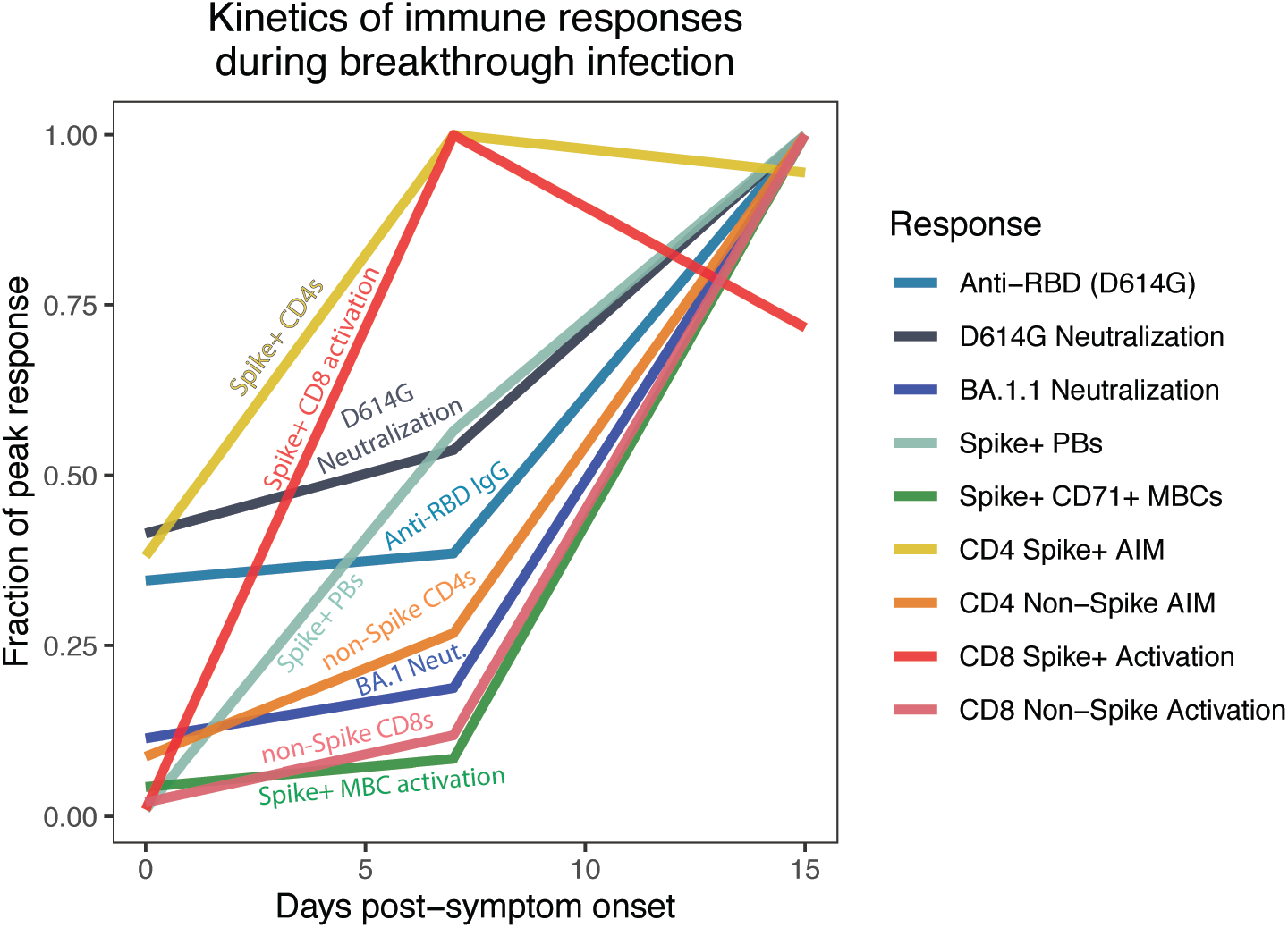
Rapid memory T cell activation and pre-existing antibodies represent the early systemic adaptive immune responses during SARS-CoV-2 breakthrough infection. Overlaid kinetics of immune responses described in Figures 1-5 depicted as the fraction of the peak response observed. The mean response value was calculated for samples at each of pre-infection baseline, day 7, and day 15.

## Discussion

Despite intense study of mRNA vaccine-induced immune memory, the dynamics with which these responses are re-engaged during SARS-CoV-2 breakthrough infection remains poorly understood. In this study, we leveraged longitudinal sampling during the symptomatic phase of infection to interrogate this key question by analyzing responses of antigen-specific antibodies, plasmablasts, memory B cells, and memory T cells during SARS-CoV-2 breakthrough infection. The findings from these studies support two overall conclusions. First, mRNA vaccination generates immunological memory that facilitates coordinated recall responses by multiple arms of the adaptive immune response during SARS-CoV-2 breakthrough infection that are more potent than primary responses observed in unvaccinated individuals. Second, though all Spike-specific responses were elevated in vaccinated individuals, not all recall responses developed with the same kinetics. Rather, re-activation of memory CD8 T cells, expansion of memory CD4 T cells, and induction of circulating plasmablasts occurred during the first week after symptom onset and preceded full activation of memory B cells in the blood or increases in the quantity and quality of neutralizing and RBD-binding antibodies. *De novo* T and B cell responses to non-Spike antigens were also not consistently detected until the second week. Despite the delayed production of new antibodies, pre-existing antibodies from prior vaccination that poorly neutralized Omicron were detectable during the first week of SARS-CoV-2 breakthrough infection. Overall, these data are consistent with the notion that pre-existing antibodies and rapidly activated memory T cells are the predominant systemic adaptive immune effectors available to limit viral replication and spread during the first week of breakthrough infection.

Antibody titers have emerged as a correlate of protection against SARS-CoV-2^1, 2^, a concept consistent with other correlates of vaccine-induced immunity^56, 57^. However, the disconnect between protection from severe disease and death versus protection from mild infection has raised questions about the potential contributions of very low levels of antibodies^46^ or memory T cells, which are predicted to limit disease severity during infection rather than preventing infection outright^58^. Moreover, whereas viral variants of concern have acquired mutations that substantially evade vaccine-induced antibody responses, Spike-specific T cell responses are substantially less impacted by these mutations^24, 25^. Our data indicate that memory CD8 T cells are robustly activated during the first week of breakthrough infection, prior to an increase in circulating antibodies. These observations, along with the emergence of *de novo* responses to non-Spike antigens during the second week, indicate that vaccine-induced humoral immunity does not sufficiently control SARS-CoV-2 replication to prevent T cell engagement or the generation of new immune responses to non-Spike antigens. Moreover, although SARS-CoV-2 has been reported to downregulate MHC class I^42, 43^, this effect did not prevent CD8 T cell priming during breakthrough infection, though it is possible that some aspects of immune evasion delayed the kinetics of these responses. Nonetheless, despite rapid induction of activation markers and Ki67, a marker of proliferation, in Spike-specific memory CD8 T cells, numerical expansion of these cells in the blood was inconsistent. One possible interpretation is that these Spike-specific memory CD8 T cells were efficiently activated and undergoing proliferative expansion, but that the expanded effector cells exit the bloodstream and accumulate in tissues at the sites of infection. Indeed, Spike-specific T cells have been detected in the nasal mucosa after breakthrough infection^59^ in humans. Further studies are warranted to interrogate the extent to which peripheral blood surveillance accurately reflects the magnitude of the antigen-specific T cell response in tissues.

The Spike-specific memory CD8 T cells that responded during breakthrough infection were predominantly CM or SCM cells, even in individuals where these populations comprised a minority of the overall Spike-specific memory CD8 T cell pool. CM and SCM cells have high proliferative potential and retain the capacity to differentiate into other effector and memory T cell subsets^60, 61^. The preferential activation and expansion of CM cells, together with the association between baseline skewing to CM lineages and overall expansion, provokes several questions that require further investigation. First, it will be interesting to identify the origin of the activated, Spike-specific CD8 CM T cells. These cells may derive from proliferation of rare pre-existing CM cells or from other subsets present at baseline, such as SCM, that may proliferate and give rise to CM cells during breakthrough infection^60^. Second, it will be informative to examine the functional quality of CD8 T cell responses skewed towards different subsets. Greater expansion in individuals with CM-skewed responses could relate to these cells having an intrinsically greater proliferative capacity, but could also result from EM-skewed responses driving accelerated antigen clearance to prevent robust expansion. Alternatively, it is possible that the activated CM and SCM in the blood reflect the residual populations of activated memory CD8 T cells that are not recruited into the tissues, while expanding effector and EM populations accumulate in tissues without detectable expansion in the blood. The heterogeneity of Spike-specific memory CD8 T cell subsets across individuals prior to breakthrough infection, together with the strong correlation between baseline frequency of Spike-specific CM and recall response upon breakthrough infection, also suggest that it will be important to define the factors that influence formation of different memory CD8 T cell subsets following mRNA vaccination. A final question is whether preferential activation of CM and SCM upon breakthrough infection relates to their ability to migrate through lymphoid tissues where dendritic cells draining sites of infection might preferentially activate these lymph node homing memory CD8 T cell populations^61^. Nevertheless, our data highlight CM cells as potentially critical responders during SARS-CoV-2 infection and highlight key questions for further exploration.

Though it is not possible to test whether memory T cell responses correlate with viral control or clinical outcomes in this study, the kinetics of activation of memory T cells suggests a role during the early stages of SARS-CoV-2 infection. Indeed, a recent study also reported activation of Spike-specific CD8 T cells during the first week of breakthrough infection, prior to increases in neutralizing antibodies, and these responses were correlated with lower peak viral load and accelerated viral clearance^62^. These authors also observed robust CD4 T cell activation during the first week of breakthrough infection, consistent with our findings of Spike-specific memory CD4 T cell expansion and activation by day 7. The current studies expand on these complementary data by identifying CD8 T cell responses to multiple HLA class I/peptide combinations and interrogating the activation and differentiation state of pre-existing and responding populations, identifying CM cells as a potentially key component of the circulating CD8 T cell recall response. Furthermore, the ability to examine antigen-specific plasmablast and memory B cell responses from the same longitudinally-sampled individuals revealed an unexpected relationship where T cell activation precedes activation of memory B cells, but occurs concomitantly with plasmablast generation. Thus, Spike-specific memory T cells are activated during breakthrough infections with accelerated kinetics compared to Spike-targeting humoral and *de novo* cellular responses to non-Spike antigens and may contribute to enhanced viral control in previously vaccinated individuals.

In contrast to the rapid activation of Spike-specific memory T cells, the maximal detection of activated Spike-specific memory B cells in the blood was delayed to the second week of infection. Memory B cells respond to re-exposure to antigen with accelerated kinetics compared to naïve B cells, seeding new germinal centers and giving rise to plasmablasts capable of rapidly producing new antibody^63, 64, 65, 66^. Though the plasmablasts detected at day 7 were likely generated from pre-existing Spike-specific memory B cells, significant increases in circulating antibody titers were not detected until day 15 post-infection. The underlying reasons for the disconnect between the circulating plasmablast and antibody responses remain unclear. The simplest explanation is that plasmablasts may need time to secrete significant quantities of antibody, as has been observed in primary responses to yellow fever or tetanus vaccination^67, 68^. This delay, however, is not observed in the setting of SARS-CoV-2 mRNA^12^ or influenza^47^ vaccination, where plasmablasts in the blood are detected with similar kinetics as new antibody. It is possible that B cell activation may be moderately impaired by a suppressive effect of pre-existing antibodies masking antigen binding sites^69, 70^. Antibody may also be bound by Spike antigen in immune complexes during the early stages of infection, precluding detection in neutralizing or binding antibody assays. Alternatively, circulating antibody may preferentially extravasate into tissues during infection as a result of local innate or adaptive immune changes, including possible vaccine-induced tissue resident memory T cells^71^. Antibodies in the lower respiratory tract are a correlate of protection in non-human primates, with anamnestic antibody detectable as early as 4 days post-challenge infection, before T cell recall responses in this model ^72, 73, 74^. These early antibody responses could feasibly be produced by memory B cells converting rapidly to antibody producing plasmablasts, as in vitro activation of human memory B cells can result in neutralizing antibody production within 4 days^14^, but may also represent exudation of pre-existing antibody^71^.

Though a rapid increase in antibodies was not observed in the blood, Spike-specific memory B cells were nonetheless potently activated during SARS-CoV-2 breakthrough infection, marked by expansion of IgG and IgA class-switched cells that could cross-bind both WT and Omicron RBD. We did not observe detectable populations of B cells specific for Omicron RBD that lacked binding to WT RBD, and the antibodies produced during the second week of infection were capable of cross-neutralizing Omicron and WT RBD. These findings suggest that memory B cells from prior vaccination dominate the response, potentially limiting *de novo* responses targeting mutated residues in variant RBDs. The possibility that B cell responses specific for the infecting variant may develop later after the resolution of infection or after multiple exposures to the variant antigen requires further investigation.

Overall, prior vaccination promoted a coordinated Spike-specific recall response during SARS-CoV-2 infection marked by increases in neutralizing antibodies, activated IgG+ and IgA+ memory B cells, and activated CD8 T cells compared to unvaccinated individuals. Multi-component recall immune responses consisting of B cells, T cells, and non-neutralizing antibodies can provide enhanced protection in animal models^75, 76^ and human H7N9 influenza infection^77^ compared to individual responses alone. Likewise, prior to widespread vaccination, coordinated immune responses characterized by simultaneous generation of SARS-CoV-2-specific antibodies, CD4 T cells, and CD8 T cells were associated with less severe COVID-19 compared to uncoordinated responses lacking detectable priming of one or more of these branches of adaptive immunity^78^. Thus, vaccination may help reduce the severity of COVID-19 during breakthrough infections by increasing the likelihood of generating a robust and coordinated adaptive immune response. Pre-existing antibodies and rapid activation of memory T cells could limit viral replication during the first week of symptoms, with memory B cell activation, rising neutralizing antibody titers, and *de novo* T cell responses to non-Spike antigens contributing to clearance of residual virus during the second week. In addition, newly produced antibodies likely serve to protect against immediate re-infection. In this way, vaccination may achieve lasting protection against severe disease by priming distinct features of adaptive immune memory to suppress viral replication throughout the course of SARS-CoV-2 breakthrough infection. Current vaccine strategies and policies, however, focus on maximizing and mainly evaluating the induction of neutralizing antibody responses. This study highlights the contribution of memory B cells and memory T cells during recall immune responses in individuals with mildly symptomatic breakthrough infections, providing a possible road map to successful vaccine-induced immunity.

## Limitations

The longitudinally-sampled Omicron breakthrough infection cohort consists of a relatively small number of individuals, mostly young, white, and all with mild COVID-19 that did not require hospitalization. The cross-sectional cohort, though larger, more diverse, and well-balanced for days post-positive test, lacks longitudinal sampling to definitively differentiate the kinetics of responses in vaccinated and unvaccinated individuals and was poorly balanced for likely infecting variant. Although the AIM assay used here demonstrates the functionality of memory CD8 T cells reactivated during breakthrough infection, it was not possible to examine this question at the level of distinct memory CD8 T cell subsets, which would require cell sorting before functional assays. The small sample size and the inability to measure viral load preclude the possibility of directly correlating immune responses with disease severity or viral replication or persistence. Furthermore, the study exclusively focuses on analysis of antibodies and immune cells circulating in peripheral blood. Though cells circulating in the blood can provide information about ongoing immune responses in tissues, as evidenced by our detection of activated T and B cells in this study, it remains possible that the kinetics of immune cell activation at the site of infection differs from what is observed in blood.

## Acknowledgments

We would like to thank the study participants for their generosity in making the study possible. We also thank the Penn Cytomics and Cell Sorting Resource Laboratory for access to instruments, Stella Park for assistance with HLA-I tetramers, and members of the Wherry lab for helpful discussions and feedback. This work was supported by grants from the NIH AI105343, AI082630, AI108545, AI155577, AI149680 (to EJW), AI152236 (to PB), HL143613 (to JRG), T32 AR076951-01 (to SAA), T32 CA009140 (to JRG and DM), T32-GM007170 (to ANS), U19AI082630 (to SEH and EJW), K08CA230157 (to ACH), funding from the Allen Institute for Immunology (to SAA, EJW), Cancer Research Institute-Mark Foundation Fellowship (to JRG), Chen Family Research Fund (to SAA), the Parker Institute for Cancer Immunotherapy (to JRG, EJW), the Penn Center for Research on Coronavirus and Other Emerging Pathogens (to PB), the University of Pennsylvania Perelman School of Medicine COVID Fund (to RRG, EJW), the Damon Runyon Clinical Investigator Award (ACH), the Doris Duke Clinical Scientist Development Award (ACH), and a philanthropic gift from Jeffrey Lurie, Joel Embiid, Josh Harris, and David Blitzer (to SEH). Work in the Wherry lab is supported by the Parker Institute for Cancer Immunotherapy. This work was also supported by NIH contract Nr. 75N9301900065 (to DW, AS). This project has been funded in part with Federal funds from the National Institute of Allergy and Infectious Diseases, National Institutes of Health, Department of Health and Human Services, under contract No. 75N93021C00015. We acknowledge the Penn Medicine BioBank (PMBB) for providing data and thank the patient-participants of Penn Medicine who consented to participate in this research program. The PMBB is approved under IRB protocol# 813913 and supported by Perelman School of Medicine at University of Pennsylvania, a gift from the Smilow family, and the National Center for Advancing Translational Sciences of the National Institutes of Health under CTSA award number UL1TR001878.

## Author Contributions

MMP and EJW conceived the study. MMP, TSJ, KAL, and JJSS carried out experiments. MMP and TJ analyzed data and wrote scripts. JCW, OK, JD, SL, MK, and CC were involved in clinical recruitment. JRG and DM provided input on statistical analyses. DM, AEB, RRG, SAA, KAL, JJSS, and ACH contributed to the methodology. MMP, DM, RRG, SAA, BF, JCW, MLM, AP, ANS, LEF, AG and SN processed peripheral blood samples. JCW and SA managed sample storage. MMP, JCW, MK, and CC managed the sample database. MMP, RRG, OK, JD, SL, and MK performed phlebotomy. JSQ, JW, MM, AG, DW, RSA, AS, and DJR provided data and materials. ARG and EJW supervised the study. All authors participated in data analysis and interpretation. MMP and EJW wrote the manuscript. All authors provided input and edits to the manuscript.

## Materials and Methods

### Human subject recruitment and sampling

Three cohorts of human subjects were recruited for this study: 1) longitudinal sampling after mRNA booster vaccination, 2) longitudinal sampling during Omicron infection of vaccinated individuals, and 3) cross-sectional collection of a single sample after a SARS-CoV-2 positive test regardless of vaccination status. Full cohort and demographic information are provided in Table S1. Additional healthy donor PBMC samples were collected with approval from the University of Pennsylvania Institutional Review Board (IRB# 845061).

1. Forty-six individuals enrolled in a longitudinal vaccine study with informed consent and approval from the University of Pennsylvania Institutional Review Board (IRB# 844642) received a 3^rd^ dose of mRNA vaccine, either Pfizer (BNT162b2) or Moderna (mRNA-1273). All participants were otherwise healthy, with no self-reported history of chronic health conditions. Peripheral blood samples (30-100mL) and clinical questionnaire data were collected at pre-vaccine baseline, day 15, and day 90 post-vaccination.
2. Eighteen individuals who had received 3-4 doses of mRNA vaccine and lacked evidence of prior SARS-CoV-2 infection were sampled longitudinally during SARS-CoV-2 infection between January and December of 2022, when Omicron and Omicron subvariants comprised the vast majority of circulating SARS-CoV-2 in the United States. For many individuals, baseline samples were acquired prior to SARS-CoV-2 infection but at least 2 months after their most recent vaccine dose. These samples are labeled as day 0 throughout the manuscript. Samples were collected during the first 15 days of infection and on day 45 based on known date of first symptom onset (IRB#851465). Prior SARS-CoV-2 infection status was determined by a combination of self-reporting and laboratory detection of pre-existing immune responses. All participants were otherwise healthy, with no self-reported history of chronic health conditions, and none were hospitalized during SARS-CoV-2 infection. Peripheral blood samples (30-100mL) and clinical questionnaire data were collected at each study visit.
3. Ninety individuals associated with the Penn Medicine Health System who tested positive for SARS-CoV-2 between April 2021 and December 2022 and were not hospitalized for COVID-19 were recruited for a single blood draw after their positive test (IRB#813913 and IRB#850981). History of SARS-CoV-2 vaccination was known for these individuals, but precise date of symptom onset was not. Fifty-nine individuals had received at least two doses of mRNA vaccine and are referred to as “vaccinated” throughout the study, while 31 were unvaccinated at the time of SARS-CoV-2 infection.

### Peripheral blood sample processing

Venous blood (30-100mL) was collected into sodium heparin tubes and plasma was separated by centrifugation at 3000rpm for 15 minutes. Heparin plasma was aliquoted and stored at −80°C for later binding and neutralizing antibody analyses. Remaining whole blood was diluted 1:1 with RPMI + 1% FBS + 2mM L-Glutamine + 100 U Penicillin/Streptomycin (R1) and layered onto SEPMATE tubes (STEMCELL Technologies) containing lymphoprep gradient (STEMCELL Technologies). After centrifugation at 1200g for 10 minutes, the PBMC fraction was harvested from SEPMATE tubes. PBMCs were then washed with R1, resuspended in ACK lysis buffer (Thermo Fisher) for 5 minutes for lysis of red blood cells, and washed again with R1. PBMCs were then filtered through a 70μm cell strainer and counted using a Countess automated cell counter (Thermo Fisher). Aliquots containing 5-20×10^6^ PBMCs were cryopreserved in 90% FBS 10% DMSO for cellular analyses.

### Measuring SARS-CoV-2 binding antibodies

SARS-CoV-2 RBD-specific binding antibody titers were tested from heparinized plasma by enzyme-linked immunosorbent assay (ELISA) as described^11, 79, 80^. Plasmids encoding the recombinant RBD were provided by F. Krammer (Mt. Sinai) and purified by nickel-nitrilotriacetic acid resin (Qiagen). On Day 0, ELISA plates (Immulon 4 HBX, Thermo Fisher Scientific) were coated with PBS or 2 μg/mL recombinant protein and stored overnight at 4°C. On Day 1, plates were washed with phosphate-buffered saline containing 0.1% Tween-20 (PBS-T) and blocked for 1 hour with PBS-T supplemented with 3% non-fat milk powder. Samples were heat-inactivated for 1 hour at 56°C and diluted in PBS-T supplemented with 1% non-fat milk powder. After washing the plates with PBS-T, 50μL of diluted sample was added to each well, and plates were incubated for 2 hours. After washing with PBS-T, 50μL of 1:5000 diluted goat anti-human IgG-HRP (Jackson ImmunoResearch Laboratories) was added to each well and plates were incubated for 1 hour. Plates were washed with PBS-T before 50μL SureBlue 3,3’,5,5’-tetramethylbenzidine substrate (KPL) was added to each well. After incubating for 5 minutes, the reaction was quenched by adding 25μL of 250 mM hydrochloric acid. Absorbance was measured with a SpectraMax 190 microplate reader (Molecular Devices) at an optical density (OD) of 450 nm. Monoclonal antibody CR3022 was included on each plate to convert WT RBD OD values into relative antibody concentrations, and endpoint titers were calculated for WT and variant RBD-binding antibodies. Plasmids to express CR3022 were provided by I. Wilson (Scripps).

### Measuring SARS-CoV-2 neutralizing antibodies

293T cells were seeded at 5×10^6^ cells per 10 cm dish, incubated for 24 hours, and transfected using calcium phosphate with 25μg of pCG1 SARS-CoV-2 S D614G delta18 or pCG1 SARS-CoV-2 S B.A.1.1 delta 18 expression plasmid encoding a codon optimized SARS-CoV-2 S gene with an 18-residue truncation in the cytoplasmic tail. The variant sequences are provided in the table below. To increase expression of the transfected DNA, fresh media containing 5mM sodium butyrate was added 12 hours post-transfection. After an additional 12 hours, the SARS-CoV-2 Spike-expressing cells were infected for 2 hours with VSV-G pseudotyped VSVΔG-RFP at an MOI of ∼1-3. Virus-containing media was removed and serum-free media was added. Media containing the SARS-CoV-2 pseudotyped VSVΔG-RFP was harvested 28-30 hours after infection, clarified by centrifugation twice at 6000xg, then aliquoted and stored at −80°C. All sera were heat-inactivated for 30 minutes at 53°C prior to use in the neutralization assay. Vero E6 cells stably expressing TMPRSS2 were seeded in 100μl at 2.5×10^4^ cells/well in a 96-well collagen-coated plate. The next day, 2-fold serially diluted serum samples were mixed with VSVΔG-RFP SARS-CoV-2 pseudotype virus (100-300 focus forming units/well) and incubated for 1 hour at 37°C. To neutralize any potential VSV-G carryover virus, a mouse anti-VSV Indiana G antibody (1E9F9), was also included at a concentration of 600 ng/ml (Absolute Antibody, Ab01402-2.0). The VeroE6 TMPRSS2 culture media was replaced with this virus-containing media, and infection was allowed to proceed for 21-22 hours. Cells were then washed and fixed with 4% paraformaldehyde before visualization on an S6 FluoroSpot Analyzer (CTL, Shaker Heights OH). Individual infected foci were enumerated and the values were compared to control wells without antibody. The focus reduction neutralization titer 50% (FRNT_50_) was calculated as the greatest serum dilution at which focus count was reduced by at least 50% relative to control cells that were infected with pseudotype virus in the absence of human serum. FRNT_50_ titers for each sample were measured in at least two technical replicates, and the geometric mean of these replicates was reported for each sample.

| Variant | Mutations in Pseudovirus Construct |
| --- | --- |
| D614G<br>(WT) | D614G |
| BA.1.1<br>(Omicron) | A67V, del69-70, T95I, G142D, del143-145, del211, L212I, ins214EPE, G339D, R346K, S371L, S373P, S375F, K417N, N440K, G446S, S477N, T478K, E484A, Q493R, G496S, Q498R, N501Y, Y505H, T547K, D614G, H655Y, N679K, P681H, N764K, D796Y, N856K, Q954H, N969K, L981F |

### SARS-CoV-2-specific plasmablast and memory B cell analyses

Antigen-specific B cells were detected using biotinylated proteins in combination with different streptavidin (SA)-fluorophore conjugates as described^11^. All reagents are listed in the **Key Resources Table**. For each sample, biotinylated proteins were multimerized with fluorescently labeled SA for 1.5 hours at 4°C at the following ratios, all of which are ∼4:1 molar ratios calculated relative to the SA-only component irrespective of fluorophore: 200ng full-length Spike protein was mixed with 20ng SA-BV421, 30ng N-terminal domain (NTD) was mixed with 12ng SA-BV786, 25ng wild-type RBD was mixed with 12.5ng SA-PE, 25ng Delta RBD was mixed with 12.5ng SA-BV711, 25ng BA.1 RBD was mixed with 12.5ng SA-PE-Cy7 and a separate 25ng BA.1 RBD was mixed with 12.5ng SA-BUV615, 25ng BA.4/5 was mixed with 12.5ng SA-APC, 50ng S2 was mixed with 12ng SA-BUV737, 50ng Nucleocapsid was mixed with 14ng SA-BV605. 12.5ng SA-PE-Cy5 was used as a decoy probe without biotinylated protein to gate out cells that non-specifically bound streptavidin. All experimental steps were performed in a 50/50 mixture of PBS + 2% FBS and Brilliant Buffer (BD Bioscience). Antigen probes were prepared individually and mixed together after multimerization with 5μM free D-biotin (Avidity LLC) to minimize potential cross-reactivity between probes. For staining, 2-10×10^6^ PBMCs per sample were thawed from cryopreservation and prepared in a 96-well round-bottom plate. Cells were first stained with Fc block (Biolegend, 1:200) and Ghost 510 Viability Dye for 15 minutes at 4°C. Cells were then washed and stained for 1 hour at 4°C with 50μL antigen probe master mix containing the above probes mixed together immediately before staining. Following incubation with antigen probe, cells were washed again and stained with anti-CD27-BUV395, anti-CD3-BUV563, anti-CD38-BUV661, anti-IgD-BV480, anti-CD19-BV750, anti-IgA-BB515, anti-CD21-PECF594, anti-IgG-AF700, and anti-CD71-APC-H7 for 30 minutes at 4°C. After surface stain, cells were washed and fixed in 1X Stabilizing Fixative (BD Biosciences) overnight at 4°C.

### Activation induced marker (AIM) stimulation assay for antigen-specific T cells

Frozen PBMC samples were thawed by warming frozen cryovials in a 37°C water bath and resuspending cells in 10mL of RPMI supplemented with 10% FBS, 2mM L-Glutamine, 100 U/mL Penicillin, and 100 μg/mL Streptomycin (R10). Cells were washed once in R10, counted using a Countess automated cell counter (Thermo Fisher), and resuspended in fresh R10 to a density of 10×10^6^ cells/mL. For each condition, 2×10^6^ cells were plated in 200μL in 96-well round-bottom plates and rested overnight in a humidifed incubator at 37°C, 5% CO_2_. After 16 hours, CD40 blocking antibody (0.5μg/mL final concentration) was added to cultures for 15 minutes prior to stimulation. Cells were then stimulated for 24 hours with costimulation (anti-human CD28/CD49d, BD Biosciences) and peptide megapools (CD4-S for all Spike-specific T cell analyses, CD4-RE for Non-Spike SARS-CoV-2-specific T cell analyses, and CD8-RE for supplemental Non-Spike SARS-CoV-2-specific CD8 T cell analyses) at a final concentration of 1 μg/mL. Peptide megapools were prepared as previously described^50, 51, 52^. Matched unstimulated samples for each donor at each timepoint were treated with costimulation alone. 20 hours post-stimulation, antibodies targeting CXCR3, CCR7, CD40L, CD107a, CXCR5, and CCR6 were added to the culture along with monensin (GolgiStop, BD Biosciences) for a four-hour stain at 37°C. After 4 hours, cells were washed in PBS supplemented with 2% FBS (FACS buffer), stained for 10 minutes at room temperature with Ghost Dye Violet 510 and Fc receptor blocking solution (Human TruStain FcX™, BioLegend), and washed once in FACS buffer. Surface staining for 30 minutes at room temperature was then performed with antibodies directed against CD4, CD8, CD45RA, CD27, CD3, CD69, CD40L, CD200, OX40, and 41BB in a 1:1 solution of FACS and Brilliant Buffer buffer. Cells were washed once in FACS buffer, fixed and permeabilizied for 30 minutes at room temperature (eBioscience™ Foxp3 / Transcription Factor Fixation/Permeabilization Concentrate and Diluent), and washed once in 1X Permeabilization Buffer prior to staining for intracellular IFN- γ overnight at 4°C. Cells were then washed once and resuspended in FACS prior to data acquisition.

All data from AIM expression assays were background-subtracted using paired unstimulated control samples. For Th1 and cTfh subsets, the AIM^+^ background frequency of non-naïve T cells was subtracted independently for each subset. AIM^+^ cells were identified from non-naïve T cell populations. AIM+ CD4 T cells were defined by dual-expression of CD200 and CD40L. AIM+ CD8 T cells were defined by dual-expression of 41BB and intracellular IFN-γ.

### Generating custom HLA-I/peptide tetramers

MHC-I tetramers were generated using the biotinylated Flex-T™ monomer UVX platform (Biolegend) for HLA-A*02:01 and HLA-A*03:01 according to manufacturer’s protocol with the modifications detailed below. Peptides were custom synthesized from Genscript, dissolved in DMSO, and stored at −20°C. For peptide exchange, all reagents were brought to 0°C on ice. Peptide stocks were diluted to 400mg/mL in cold PBS, and 20μL of diluted peptide was mixed with 20μL of Flex-T™ monomer UVX (200μg/ml) by pipetting up and down in a 96-well V bottom plate. The plate was sealed and centrifuged at 2000rpm for 2 minutes at 4°C to collect the liquid down. The seal was removed, and the plate was illuminated with 366nm UV light at 4°C for 30 minutes at a distance of 2-5cm from the UV lamp. After 30 minutes exposure to UV light, the plate was covered and incubated for 30 minutes at 37°C in the dark.

To generate tetramers, the plate was spun at 2000rpm for 2 min at 4°C, at which point 30μL from the top of each well was carefully transferred into PCR tube strips. A total of 0.66μg of fluorophore-conjugated streptavidin was added, and tetramers were incubated on ice in the dark for 30 minutes. Tetramers were stored in the dark at 4°C for up to 2 months. The fluorophore pairs for peptides from each SARS-CoV-2 protein were as follows: Spike peptides were multimerized BB515 and PE, Nucleocapsid peptides were multimerized with BB515 and BV650, ORF3a peptides were multimerized with BB515 and BV786, and Replicase peptides were multimerized with BV650 and PE.

### HLA-I/peptide tetramer staining

Cryopreserved PBMCs were thawed as for the AIM assay, counted, and stained at a concentration of 2.5×10^6^ cells per well in a 96-well round bottom plate. All incubations were performed in the dark, and staining was performed in 50μL volume. All stains and washes were performed in FACS buffer (PBS + 2% FBS) unless otherwise noted. HLA-I/peptide tetramer staining solution was prepared by mixing all tetramers into a solution of 1:1 FACS buffer and Brilliant Buffer containing 0.2% milk and 5μM free D-biotin (Avidity LLC) immediately prior to staining. Individual tetramer reagents conjugated to each of two fluorophores were added for each peptide antigen, for a total of 44 tetramers (22 unique peptides) for HLA-A*02:01 staining and 14 tetramers (7 unique peptides) for HLA-A*03:01. To avoid aggregates, tetramer stocks were briefly centrifuged prior to adding to the staining solution. Cells were pelleted by centrifugation at 1600rpm for 3 minutes, supernatant was removed, and cells were resuspend in 50μL of HLA-I/peptide tetramer staining solution. After incubating for 20 minutes at room temperature, 150mL of FACS buffer was added, samples were spun at 1600rpm for 3 minutes, and the supernatant was removed (“wash”). Samples were resuspended in 50μL of Ghost viability dye and Fc blocking solution, incubated for 10 minutes at room temperature, washed, and resuspended in 50μL of surface antibody cocktail in 1:1 FACS buffer and Brilliant Buffer containing antibodies targeting CD45RA, CD8, CD39, CX3CR1, CD27, CD3, PD-1, CD49d, CD14 (dump), CD16 (dump), CD19 (dump), CD41a (dump), HLA-DR, CD38, CD127, CD95, CD4, CCR7, and KLRG1. After incubating for 30 minutes at room temperature, cells were washed and resuspended in 100μL of Fixation/Permeabilization solution (eBioscience™ Foxp3 / Transcription Factor Fixation/Permeabilization Concentrate and Diluent), incubated 30 minutes at room temperature, and washed with 100μL of 1X Perm Buffer. Cells were then pelleted at 2000rpm for 5 minutes, the supernatant was removed, and cells were resuspended in 50μL of intracellular staining solution containing antibodies directed against Granzyme B, T-bet, TCF-1, and Ki67 in 1X Perm Buffer and incubated at 4°C overnight. The next day, cells were washed in 1X Perm Buffer, resuspended in FACS buffer, filtered through a cell strainer, and analyzed on a BD Symphony A5 cytometer.

### Flow cytometry

Samples were acquired on a BD Symphony A5 instrument. Standardized SPHERO rainbow beads (Spherotech) were used to track and adjust photomultiplier tubes over time. UltraComp eBeads (Thermo Fisher) were used for compensation. Up to 5×10^6^ cells were acquired per sample. Data were analyzed using FlowJo v10 (BD Bioscience) or OMIQ.

## QUANTIFICATION AND STATISTICAL ANALYSIS

### High dimensional analysis and statistics

All data were analyzed using custom scripts in R and visualized using RStudio. Statistical tests are indicated in the corresponding figure legends. All tests were performed two-sided with a nominal significance threshold of p < 0.05. Benjamini-Hochberg (BH) correction was performed in all cases of multiple comparisons. Unpaired tests were used for comparisons between timepoints unless otherwise indicated as some participants were missing samples from individual timepoints. * indicates p < 0.05, ** indicates p < 0.01, *** indicates p < 0.001, **** indicates p < 0.0001. Source code and data files are available upon request from the authors.

