## Supplemental Materials for "Prior vaccination enhances immune responses during SARS-CoV-2 breakthrough infection with early activation of memory T cells followed by production of potent neutralizing antibodies"

**Supplementary Information**

**Contents**

Table S1: Cohort Summary
Figure S1
Figure S2
Figure S3
Figure S4
Figure S5
Key Resources Table
Penn Medicine BioBank Banner Author List and Contribution Statements

### Table S1: Cohort Summary

| Characteristics | | Longitudinal Omicron Cohort | Cross-sectional Unvaccinated | Cross-sectional Vaccinated |
|---|---|---|---|---|
| Total | Number | 18 | 31 | 59 |
| Age | Average (Years) | 35.6 | 38.5 | 43.6 |
| | Range (Years) | 23-60 | 23-64 | 23-73 |
| | 20-30 | 7 (39%) | 9 (29%) | 12 (20%) |
| | 30-40 | 6 (33%) | 10 (32%) | 15 (25%) |
| | 40-50 | 2 (11%) | 5 (16%) | 8 (14%) |
| | 50+ | 3 (17%) | 7 (23%) | 24 (41%) |
| Sex | Male | 8 (44%) | 13 (42%) | 25 (42%) |
| | Female | 10 (56%) | 18 (58%) | 34 (58%) |
| Race/Ethnicity | White - Non-Hispanic/Latino | 14 (77%) | 1 (3%) | 25 (42%) |
| | White - Hispanic/Latino | 1 (6%) | 0 | 2 (3%) |
| | Asian | 3 (17%) | 0 | 6 (10%) |
| | Black | 0 | 30 (97%) | 24 (41%) |
| | Native | 0 | 0 | 0 |
| | Other | 0 | 0 | 2 (3%) |
| HLA Type | HLA-A*02:01 | 9 (50%) | 11 (35%) | 19 (32%) |
| | HLA-A*03:01 | 3 (17%) | 3 (10%) | 8 (14%) |
| Vaccine doses | 2 | 0 | 0 | At least 2 doses: 59 (100%) |
| | 3 | 16 (89%) | 0 | |
| | 4 | 2 (11%) | 0 | |
| Dominant Variant | Alpha | 0 | 15 (48%) | 3 (5%) |
| | Delta | 0 | 14 (45%) | 28 (47%) |
| | Omicron | 18 (100%) | 2 (6%) | 28 (47%) |
| Days post-test | Mean (IQR) | NA | 8.5 (6 - 10.5) | 9.7 (7 - 13) |

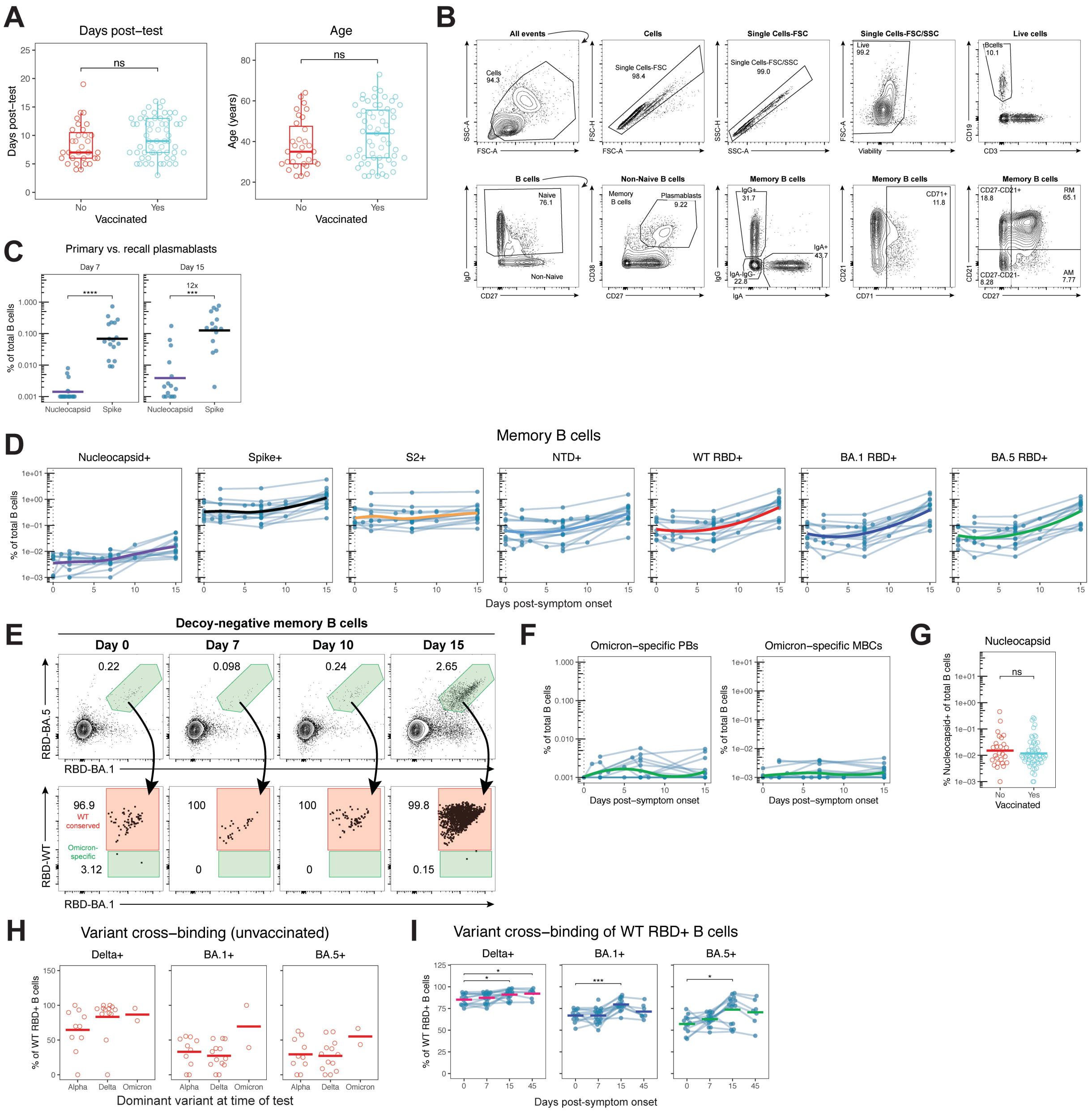

**Fig. S1: Supplement for Figures 1-2. A)** Days between positive test and sample collection (left) and age (right) for the vaccinated and unvaccinated individuals in the cross-sectional cohort. **B)** Representative flow cytometric gating strategy for identifying plasmablasts, memory B cells, and memory B cell phenotypes. RM = resting memory, AM = activated memory. **C)** Frequency of Nucleocapsid and Spike-specific plasmablasts during Omicron breakthrough infection. **D)** Flow cytometric data depicting the frequency of total B cells that are memory B cells binding Spike or Nucleocapsid antigens during Omicron breakthrough infection. **E)** Representative flow cytometric gating strategy for identifying B cells specifically binding Omicron subvariant RBDs with or without conserved binding to WT RBD, gated directly from Decoy-negative B cells without prior gating on cells binding the WT Spike probe. **F)** Frequency of Omicron-specific plasmablasts (left) and memory B cells (right), defined as cells binding to both the BA.1 RBD and BA.5 RBD probes without binding the WT RBD probe. **G)** Frequency of total B cells that are Nucleocapsid-specific memory B cells during SARS-CoV-2 infection of previously vaccinated and unvaccinated individuals. **H)** Variant cross-binding of WT RBD-binding memory B cells during SARS-CoV-2 infection of unvaccinated individuals, divided by dominant variant at the time of positive test. **I)** Variant cross-binding of WT RBD-binding memory B cells during Omicron breakthrough infection. Statistics calculated using Wilcoxon test with Benjamini-Hochberg correction for multiple comparisons.

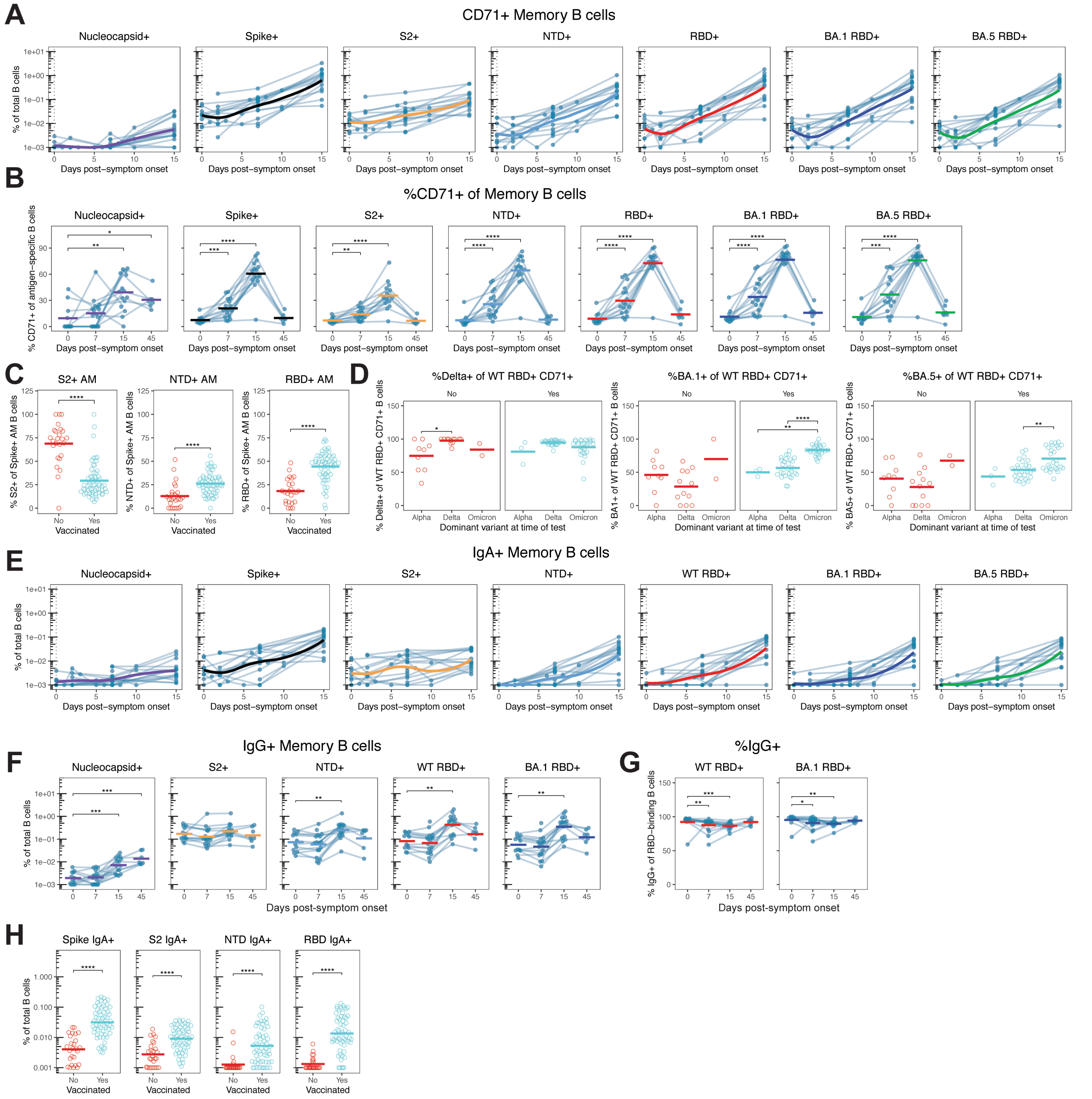

**Fig. S2: Supplement for Figure 2. A)** Frequency of total B cells that are activated (CD71+) memory B cells and bind Spike or Nucleocapsid antigens during Omicron breakthrough infection. **B)** Percent activation (CD71+) of antigen-specific memory B cells. **C)** Percent of activated memory (AM) Spike-binding memory B cells targeting the S2, NTD, or RBD domains during SARS-CoV-2 infection, used as an alternative to CD71 but supporting the same conclusions. **D)** Percent of WT RBD-binding CD71+ memory B cells that cross bind the indicated variant RBDs during SARS-CoV-2 breakthrough infection separated by vaccination status and the dominant circulating variant at the time of positive test. **E)** Frequency of total B cells that are IgA+ memory B cells and bind Spike or Nucleocapsid antigens during Omicron breakthrough infection. **F)** Frequency of total B cells that are IgG+ memory B cells and bind Spike or Nucleocapsid antigens during Omicron breakthrough infection. **G)** Percent of WT and BA.1 RBD-binding B cells that are IgG+ during breakthrough infection. **H)** Frequency of total B cells that are IgA+ memory B cells and specific for the indicated domains of Spike during SARS-CoV-2 infection in previously vaccinated or unvaccinated individuals. Statistics calculated using Wilcoxon test with Benjamini-Hochberg correction for multiple comparisons.

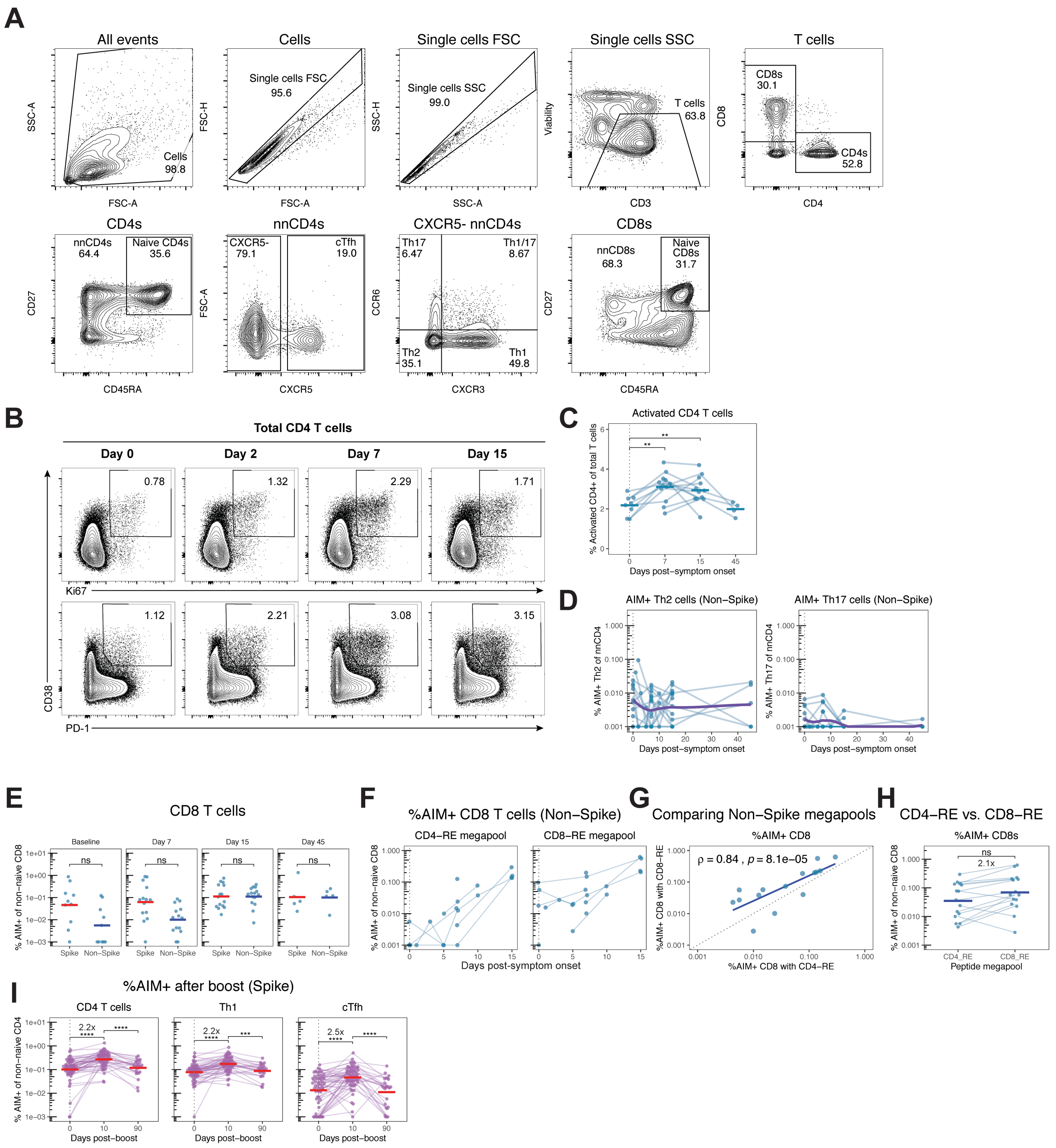

**Fig. S3: Supplement for Figure 3. A)** Representative flow cytometric gating strategy for identifying non-naïve CD4 and CD8 T cells and CD4 T cell subsets. **B)** Representative gating showing the kinetics of bulk CD4 T cell activation during Omicron breakthrough infection. **C)** Bulk activation of CD4 T cells during Omicron breakthrough infection calculated as percent of total T cells that are CD4 T cells expressing at least 2 of 5 activation markers as in Fig. S4C. **D)** Percent of non-naïve CD4 T cells that are Spike-specific Th2 (CXCR5-, CXCR3-, CCR6-) or Th17 (CXCR5-, CXCR3-, CCR6+) cells. **E)** Comparison of total Spike- and Non-Spike-specific CD8 T cell responses during Omicron infection. **F)** Comparison of non-Spike-specific CD8 T cell responses detected using CD4-RE and CD8-RE peptide megapools on a subset of samples collected during Omicron breakthrough infection. **G-H)** Correlation (G) and comparison (H) of CD8 T cell responses detected from paired samples stimulated with CD4-RE and CD8-RE, excluding samples with undetectable responses to either megapool. Dotted line indicates a hypothetical 1:1 relationship in the detection of CD8 T cells specific for non-Spike epitopes using each megapool. **I)** Frequency of Spike specific CD4 T cells, Th1, and cTfh before and after a 3rd dose of mRNA vaccine (cohort from Fig. 1A). Statistics calculated using Wilcoxon test with Benjamini-Hochberg correction for multiple comparisons.

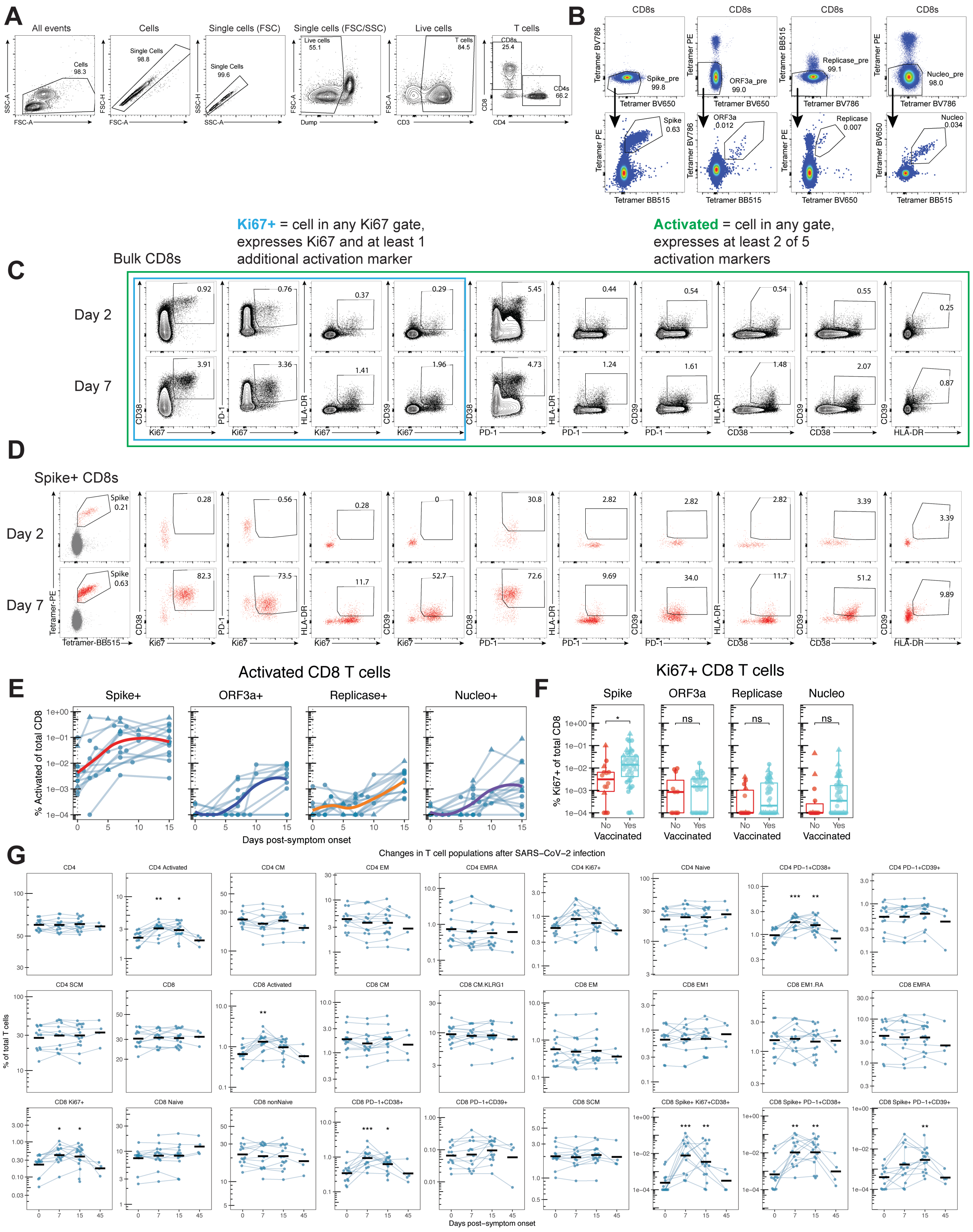

**Fig. S4: Supplement for Figure 4. A)** Representative flow cytometric gating strategy for identifying total CD8 T cells for MHC-I tetramer staining and phenotyping. **B)** Representative flow cytometric gating strategy for identifying MHC-I tetramer-binding cells from total CD8 T cells. Cells are pre-gated to not fluoresce in the two mismatched tetramer channels before identifying antigen-specific cells based on dual-fluorescence as depicted in Fig. 4A. **C-D)** Representative gating of activated C) bulk and D) Spike-specific CD8 T cells on day 2 and day 7 of breakthrough infection. Cells defined as Ki67+ fall in any gate co-expressing Ki67 and another activation marker (blue), and cells defined as activated fall in any gate co-expressing two activation markers (green). **E)** Percent of total CD8 T cells that are antigen-specific and express at least two activation markers, supporting Figure 4H with an alternative method for defining activated cells. **F)** Percent of total CD8 T cells that are antigen-specific and express Ki67 and at least one other activation marker during SARS-CoV-2 infection in previously vaccinated or unvaccinated individuals, supporting Figure 4J with an alternative method for defining activated cells. **G)** Percent of total T cells attributed to various bulk CD4 or CD8 T cell populations or Spike-specific CD8 T cell populations during Omicron breakthrough infection, highlighting transient activation of CD4 and CD8 T cells but no lasting alterations to the T cell landscape by 45 days after mild SARS-CoV-2 infection. Statistics calculated using Wilcoxon test with Benjamini-Hochberg correction for multiple comparisons. Statistics without brackets are in comparison to day 0.

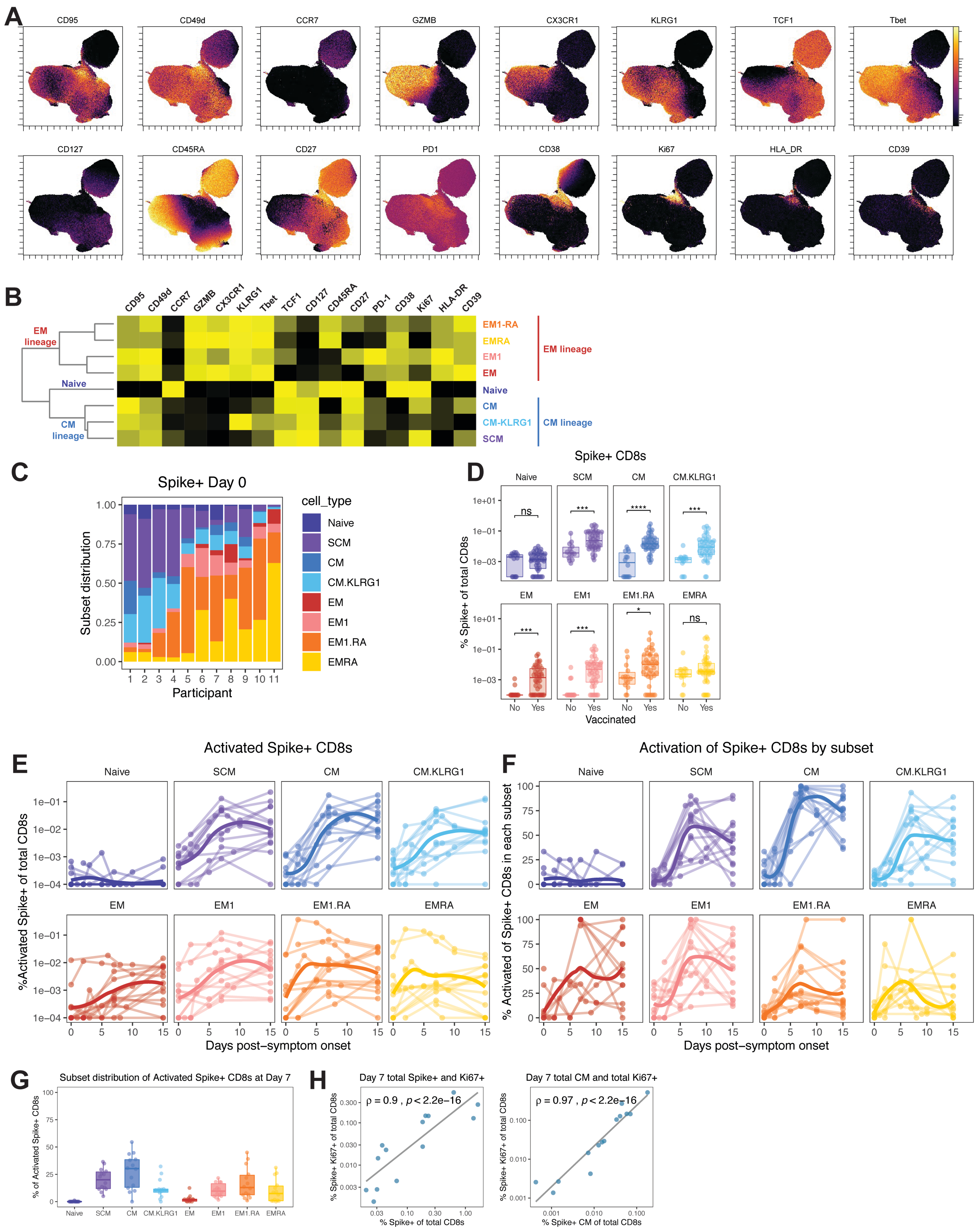

**Fig. S5: Supplement for Figure 5. A)** Expression in UMAP space of each of the 16 input parameters used to generate the UMAP in Fig. 5B. **B)** Heat map depicting average expression of each marker in the manually gated populations as defined in Fig. 5A. The dendrogram at the left clusters populations based on the similarity of their average expression across all 16 parameters.

**C)** Subset distribution of Spike-specific CD8 T cells at pre-infection baseline for eleven subjects with paired measurements at baseline and day 7, as in Fig. 5H. **D)** The abundance of Spike-specific CD8 T cells in each subset during SARS-CoV-2 infection of previously vaccinated or unvaccinated individuals, calculated as the percent of total CD8 T cells. **E)** The magnitude of the activated Spike-specific CD8 T cell response in each subset during breakthrough infection as a percent of total CD8 T cells, where activated cells express at least two activation markers. Supports Fig. 5E using a different method for defining activated cells. **F)** The percent activation of Spike-specific CD8 T cells in each subset, where activated cells express at least two activation markers. Supports Fig. 5G using a different method for defining activated cells. **G)** Summary data for the subset distribution of activated Spike-specific CD8 T cells at day 7 post-symptom onset. Supports Fig. 5I using a different method for defining activated cells. **H)** Correlations within Spike-specific CD8 T cell responses. Left) Day 7 total Spike-specific CD8 T cells and Day 7 total Ki67+ Spike-specific CD8 T cells. Right) Day 7 total Spike-specific CM cells and Day 7 total Ki67+ Spike-specific CD8 T cells. Statistics calculated using Wilcoxon test with Benjamini-Hochberg correction for multiple comparisons.

### KEY RESOURCES TABLE

| REAGENT or RESOURCE | SOURCE | IDENTIFIER |
|---|---|---|
| Antibodies | | |
| BUV563 anti-CD3 | BD Biosciences | Cat#748569 |
| BV750 anti-CD19 | Biolegend | Cat#302262 |
| BUV805 anti-CD20 | BD Biosciences | Cat#612905 |
| BUV395 anti-CD27 | BD Biosciences | Cat#563815 |
| BUV661 anti-CD38 | BD Biosciences | Cat#612969 |
| APC-H7 anti-CD71 | BD Biosciences | Cat#563671 |
| AF700 anti-CD11c | Biolegend | Cat#337220 |
| FITC anti-IgA | Miltenyi | Cat#130-113-475 |
| BV480 anti-IgD | BD Biosciences | Cat#566138 |
| PE/Dazzle 594 anti-CD21 | Biolegend | Cat#354922 |
| PE-Cy7 anti-IgG | Biolegend | Cat#410722 |
| PerCP/Cy5.5 anti-IgM | Biolegend | Cat#314512 |
| mouse anti-VSV Indiana G, 1E9F9 | Absolute Antibody | Cat#Ab01402-2.0 |
| goat anti-human IgG-HRP | Jackson ImmunoResearch Laboratories | RRID: AB_2337596 |
| BUV395 CD4 | BD Biosciences | Cat#563550 |
| BUV496 CD8 | BD Biosciences | Cat#612943 |
| BUV615 CD45RA | BD Biosciences | Cat#751555 |
| BUV737 CD27 | BD Biosciences | Cat#612829 |
| BUV805 CD3 | BD Biosciences | Cat#612896 |
| BV421 CXCR3 | Biolegend | Cat#353716 |
| BV650 CCR7 | Biolegend | Cat#353234 |
| BV605 CD69 | Biolegend | Cat#310938 |
| BV711 CD40L | Biolegend | Cat#310838 |
| BV785 CD107a | Biolegend | Cat#328644 |
| FITC IFNy | Biolegend | Cat#502515 |
| PE CD200 | Biolegend | Cat#399804 |
| PE-Cy7 OX40 | Biolegend | Cat#350012 |
| AF647 41BB | Biolegend | Cat#309810 |
| APC-R700 CXCR5 | BD Biosciences | Cat#565191 |
| APC-Cy7 CCR6 | Biolegend | Cat#353432 |
| BB700 CD127 (anti-human) | BD | 566398 |
| AF647 TCF1 (anti-human) | CellSignaling | 6709S |
| AF700 CCR7 (anti-human) | Biolegend | 353244 |
| APC/Fire750 KLRG1 (anti-human) | Biolegend | 367718 |
| BUV395 CD45RA (anti-human) | BD | 740298 |
| BUV496 CD8 (anti-human) | BD | 612942 |
| BUV661 CX3CR1 (anti-human) | BD | 750690 |
| BUV737 CD27 (anti-human) | BD | 612830 |
| BUV805 CD3 (anti-human) | BD | 612896 |
| BV421 PD-1 (anti-human) | Biolegend | 329920 |
| BV480 CD49d (anti-human) | BD | 566134 |
| V500 CD14 (anti-human) | BD | 561391 |
| V500 CD16 (anti-human) | BD | 561394 |

| | | |
|---|---|---|
| V500 CD19 (anti-human) | BD | 561121 |
| BV510 CD41a (anti-human) | BD | 563250 |
| BV605 HLA-DR (anti-human) | Biolegend | 307640 |
| BV711 Ki67 (anti-human) | Biolegend | 350516 |
| BV750 CD38 (anti-human) | BD | 746909 |
| PE-Cy5 CD95 (anti-human) | BD | 561977 |
| PE-Cy5.5 CD4 (anti-human) | Invitrogen | 35-0047-42 |
| PE-Cy7 T-bet (anti-human) | Biolegend | 644824 |
| PE-TexasRed Granzyme B (anti-human) | Invitrogen | GRB17 |
| BUV563 CD39 (anti-human) | BD | 748473 |
| **Bacterial and virus strains** | | |
| SARS-CoV-2 VSV pseudotypes | Generated for this paper | |
| **Biological samples** | | |
| Human peripheral blood samples | Collected at the University of Pennsylvania | N/A |
| **Chemicals, peptides, and recombinant proteins** | | |
| SARS-CoV-2 Biotyinlated Full Length Spike | R&D Systems | Cat#AVI10549-050 |
| HA(ΔTM)(A/Brisbane/02/2018)(H1N1) | Immune Tech | Cat#IT-003-00110ΔTMp |
| HA(ΔTM)(B/Colorado/06/2017) | Immune Tech | Cat#IT-003-B21ΔTMp |
| SARS-CoV-2 Biotyinlated RBD (WT) | Acro Biosystems | Cat#SPD-C82E9-25ug |
| SARS-CoV-2 Biotyinlated RBD (Delta) | Acro Biosystems | Cat#SPD-C82E5-25ug |
| SARS-CoV-2 Biotyinlated RBD (Omicron BA.5) | Acro Biosystems | Cat#SPD-C82Ew-25ug |
| SARS-CoV-2 Biotyinlated RBD (Omicron BA.1) | Acro Biosystems | Cat#SPD-C82E4-25ug |
| SARS-CoV-2 Biotyinlated N-Terminal Domain | Sino Biological | Cat#40591-V49H-B |
| SARS-CoV-2 Biotyinlated S2 | Acro Biosystems | Cat#S2N-C52E8-25ug |
| SARS-CoV-2 Biotyinlated Nucleocapsid | R&D Systems | Cat#BT10474-050 |
| SARS-CoV-2 (BA.1.1) Spike RBD Protein (His Tag) | Sino Biological | Cat#40592-V08H129 |
| SARS-CoV-2 (BA.4/BA.5) Spike RBD Protein (His Tag) | Sino Biological | Cat#40592-V08H130 |
| BV421 Streptavidin | Biolegend | Cat#405226 |
| BV605 Streptavidin | Biolegend | Cat#405229 |
| BV711 Streptavidin | BD Biosciences | Cat#563262 |
| BV786 Streptavidin | BD Biosciences | Cat#563858 |
| BUV615 Streptavidin | BD Biosciences | Cat#613013 |
| BUV737 Streptavidin | BD Biosciences | Cat#612775 |
| BB515 Streptavidin | BD Biosciences | Cat#564453 |
| PE Streptavidin | Biolegend | Cat#405203 |
| PE-Cy7 Streptavidin | Biolegend | Cat#405206 |
| APC Streptavidin | Biolegend | Cat#405207 |
| Ghost Viability Dye Violet 510 | Tonbo | Cat#13-0870-T100 |
| Human TruStain FcX (Fc Receptor Blocking Solution) | Biolegend | Cat#422302 |
| EZ-Link Micro NHS-PEG4 Biotinylation Kit | Thermo Fisher | Cat#21955 |
| Zeba Spin Desalting Columns 7K MWCO | Thermo Fisher | Cat#89894 |
| CD4-S peptide megapool | Synthetic Biomolecules (aka A&A) | http://www.syntheticbiomolecules.com/ |
| CD4-RE peptide megapool | Synthetic Biomolecules (aka A&A) | http://www.syntheticbiomolecules.com/ |
| CD8-RE peptide megapool | Synthetic Biomolecules (aka A&A) | http://www.syntheticbiomolecules.com/ |

| | | |
|---|---|---|
| GolgiStop (Containing Monensin) | BD Biosciences | Cat#51-2092K7 |
| CD40 Antibody, anti-human, pure-functional grade | Miltenyi Biotech | Cat#130-094-133 |
| Anti-Human CD28/CD49d Purified | BD Biosciences | Cat#347690 |
| Foxp3 / Transcription Factor Fixation/Permeabilization Concentrate and Diluent | eBioscience | Cat#00-5521-00 |
| ALWEIQQVV | Genscript | Custom synthesis |
| FLAHIQWMV | Genscript | Custom synthesis |
| FLLNKEMYL | Genscript | Custom synthesis |
| KLWAQCVQL | Genscript | Custom synthesis |
| SMWALIISV | Genscript | Custom synthesis |
| YLATALLTL | Genscript | Custom synthesis |
| LALLLLDRL | Genscript | Custom synthesis |
| GMSRIGMEV | Genscript | Custom synthesis |
| LLLDRLNQL | Genscript | Custom synthesis |
| ALSKGVHFV | Genscript | Custom synthesis |
| LLYDANYFL | Genscript | Custom synthesis |
| ALNTLVKQL | Genscript | Custom synthesis |
| KIADYNYKL | Genscript | Custom synthesis |
| RLQSLQTYV | Genscript | Custom synthesis |
| SIIAYTMSL | Genscript | Custom synthesis |
| VVFLHVTYV | Genscript | Custom synthesis |
| FIAGLIAIV | Genscript | Custom synthesis |
| NLNESLIDL | Genscript | Custom synthesis |
| RLNEVAKNL | Genscript | Custom synthesis |
| TLDSKTQSL | Genscript | Custom synthesis |
| VLNDILSRL | Genscript | Custom synthesis |
| YLQPRTFLL | Genscript | Custom synthesis |
| KLFAAETLK | Genscript | Custom synthesis |
| KLFDRYFKY | Genscript | Custom synthesis |
| KTIQPRVEK | Genscript | Custom synthesis |
| VTNNTFTLK | Genscript | Custom synthesis |
| KTFPPTEPK | Genscript | Custom synthesis |
| GVYFASTEK | Genscript | Custom synthesis |
| KCYGVSPTK | Genscript | Custom synthesis |
| Flex-T™ HLA-A*02:01 Monomer UVX | BioLegend | Cat#280004 |
| Flex-T™ HLA-A*03:01 Monomer UVX | BioLegend | Cat#280005 |
| **Experimental models: Cell lines** | | |
| 293T | ATCC | RRID: CVCL_0063 |
| VeroE6/TMPRSS | Stefan Pohlman | Anderson et al., 2021 |
| **Recombinant DNA** | | |
| Plasmid: pCAGGS SARS-CoV-2 spike | Florian Krammer | Amanat et al. (2020) |
| Plasmid: pCAGGS SARS-CoV-2 RBD | Florian Krammer | Amanat et al. (2020) |
| Plasmid: pCG1 SARS-CoV-2 D614G delta18 | Paul Bates Lab | N/A |
| Plasmid: pCG1 SARS-CoV-2 B.A.1.1 delta18 | Paul Bates Lab | N/A |

**Penn Medicine BioBank Banner Author List and Contribution Statements**

**PMBB Leadership Team**
Daniel J. Rader, M.D., Marylyn D. Ritchie, Ph.D., Michael D. Feldman M.D.

Contribution: All authors contributed to securing funding, study design and oversight. All authors reviewed the final version of the manuscript.

**Patient Recruitment and Regulatory Oversight**
JoEllen Weaver, Nawar Naseer, Ph.D., M.P.H., Afiya Poindexter, Ashlei Brock, Khadijah Hu-Sain, Yi-An Ko

Contributions: JW manages patient recruitment and regulatory oversight of study. NN manages participant engagement, assists with regulatory oversight, and researcher access. AP, AB, KH, YK perform recruitment and enrollment of study participants.

**Lab Operations**
JoEllen Weaver, Meghan Livingstone, Fred Vadivieso, Ashley Kloter, Stephanie DerOhannessian, Teo Tran, Linda Morrel, Ned Haubein, Joseph Dunn

Contribution: JW, ML, FV, SD conduct oversight of lab operations. ML, FV, AK, SD, TT, LM perform sample processing. NH, JD are responsible for sample tracking and the laboratory information management system.

**Clinical Informatics**
Anurag Verma, Ph.D., Colleen Morse, M.S., Marjorie Risman, M.S., Renae Judy, B.S.

Contribution: All authors contributed to the development and validation of clinical phenotypes used to identify study subjects and (when applicable) controls.

**Genome Informatics**
Anurag Verma Ph.D., Shefali S. Verma, Ph.D., Yuki Bradford, M.S., Scott Dudek, M.S., Theodore Drivas, M.D., PH.D.

Contribution: A.V., S.S.V. are responsible for the analysis, design, and infrastructure needed to quality control genotype and exome data. Y.B. performs the analysis. T.D. and A.V. provides variant and gene annotations and their functional interpretation of variants.
